# Simulating Thermoresponsive Behavior of Disordered Proteins with Temperature-Dependent Coarse-Grained Potentials Derived from Hydration Free Energies

**DOI:** 10.64898/2026.01.30.702805

**Authors:** Fangke Chen, Xiangze Zeng

## Abstract

Thermoresponsive phase transitions of intrinsically disordered proteins (IDPs), including both upper critical solution temperature (UCST) and lower critical solution temperature (LCST) transitions, are widely observed in natural and synthetic sequences. However, most existing coarse-grained (CG) models employ temperature-independent interactions and fail to capture solvation-driven LCST behavior. To address this, we introduce **TEA** (**T**emperature-dependent **E**nergetics derived from hydr**A**tion free energies), a physics-based framework that augments CG models with temperature-dependent interactions derived from hydration thermodynamics. Leveraging extensive all-atom molecular dynamics simulations, we demonstrate that inter-residue interaction strengths quantified by excess free energies correlate linearly with hydration free energies. This correlation, combined with a validated combination rule for heterotypic interactions, allows the derivation of temperature-dependent potentials across continuous temperature space. We map these atomistic energetics to CG potentials via a single global scaling parameter, ensuring minimal overfitting and high transferability. The TEA-augmented CG models robustly distinguish UCST- and LCST-type sequences, successfully identify experimentally reported outliers, and accurately reproduce LCST-type single-chain compaction trends and phase diagrams of multiple disordered proteins. Collectively, our work provides a transferable and physically interpretable framework that bridges atomistic hydration thermodynamics and phase behavior of IDPs, enabling the simulation of thermoresponsive sequences with minimal phenomenological fitting.

## Introduction

Biomolecular phase separation has emerged as a fundamental principle for understanding various cellular processes [1-4]. Recent studies have demonstrated the crucial roles of intrinsically disordered proteins or regions (IDPs) as drivers or regulators of cellular phase separation, owning to their multivalent interactions [5-9]. Unlike folded proteins, IDPs lack stable tertiary structures and instead sample a heterogeneous ensemble of conformations dictated by their low fraction of hydrophobic residues. This structural flexibility enables IDPs to function as flexible linkers between folded domains, molecular recognition motifs for protein binding, and molecular assemblers promoting higher-order structure formation [10]. Consequently, elucidating the conformational ensembles and phase behavior of IDPs is crucial for understanding the molecular basis of their biological functions.

Molecular simulations, including molecular dynamics (MD) and Monte-Carlo methods, are essential tools for probing protein dynamics at high spatiotemporal resolution [11-13]. However, atomistic simulations of IDPs face two major challenges. First, widely used protein force fields are typically parameterized for folded proteins, leading to overly collapsed states when applied to IDPs [14-16]. To mitigate this issue, various modifications have been made to these force fields [17-23]. For example, Best et al scaled the protein-water interactions in the Amber ff03 force fields to prevent overly compaction [17]. Besides modifying protein-water interactions, Robustelli et al refined torsion parameters in Amber 99sb-ILDN force fields to balance helix and coil populations [18]. Huang et al introduced an improved CMAP backbone potential in CHARMM36m force fields [19]. The second challenge is the high computational cost for simulating extended IDP conformations, requiring large simulation boxes with explicit water molecules that significantly slows down simulation speed. An approach to address this challenge is using implicit solvent model. For instance, the ABSINTH all-atom model has been developed and parameterized for IDPs [24]. While these all-atom force fields have achieved some success in capturing the single chain conformations of IDPs, simulating the phase separation involving at least hundreds of IDPs remains computationally challenging at the atomistic level.

To bridge these spatiotemporal scales, coarse-grained (CG) simulations have been widely adopted for studying IDP phase separation [25-34]. For instance, Choi et al developed the phenomenological lattice model LaSSI, which can capture the essential underlying physical principles of protein phase separation [25, 35]. Dignon et al developed the one-bead-per-residue CG model HPS based on residue hydrophobicity scale [27]. Tesei et al further improved the parameters of HPS model using a data-driven approach and proposed the Calvados model. Similarly, Joseph et al developed the mpipi model using both atomistic simulation and bioinformatic data [28]. These models successfully capture phase separation with upper critical solution temperature (UCST), where phase separation is driven by favorable enthalpy upon cooling.

However, these models fail to describe phase separation with lower critical solution temperature (LCST), where phase separation occurs upon heating. This limitation arises from the temperature-independent inter-residue energies in these implicit solvent models. One physical origin of LCST-type phase behavior is the solvation effect, where residues become less favorably solvated at higher temperatures [36, 37]. Meanwhile, Wadworth et al showed that ion effect is another driving force for LCST-type phase separation [38]. Specifically, they found that more Mg^2+^ ions bridges RNA molecules by binding to multiple phosphate groups at elevated temperatures, which drives phase separation. In both cases, the effective interactions between solute molecules are increased with temperatures. Since these CG models employ implicit solvents, they cannot fully capture these solvation effect or ion effects.

It has been shown that the all-atom implicit solvent model ABSINTH has successfully reproduced the LCST-type phase behavior by including temperature-dependent solvation term in the Hamiltonian [36]. Wuttke et al incorporated experimentally inferred temperature-dependent solvation free energies to ABSINTH and observed the LCST-type single-chain coil-to-globule transitions for five IDPs [36]. Building on this, we previously calculated temperature-dependent hydration free energies for the backbone and sidechain analogues from all-atom simulations with the polarizable force fields AMOEBA [39] and developed the ABINTH-T model [37]. Combined with a generic algorithm, we successfully designed synthetic IDPs with LCST-type phase behavior, which have been validated by preliminary experimental measurements [37].

Several attempts have been made to incorporate temperature-dependent potentials to coarse-grained models [40, 41]. Based on the HPS model, Dignon et al used a parabolic function to describe the temperature dependent interaction parameter *λ* [40]. The final parameters were obtained by fitting the single-chain coil-to-globule transitions of five LCST-type IDPs from Wuttke’s experimental study [36]. Recently, Chakravarti and Joseph used a data-driven approach to fit the coarse-grained temperature dependent interactions [41]. Specifically, built on the insights of Wuttke [36] and the ABSINTH-T model [37], they used the function form of hydration free energies to fit the temperature-dependent interaction strengths in mpipi model. These efforts support that integrating the temperature-dependent potentials to the coarse-grained models can capture the LCST-type phase behavior. However, while experimental data for LCST-type phase behavior is limited, it is still challenging obtaining consistent temperature-dependent energetics across residue chemistries, constructing heterotypic interactions without an explosion of fitted parameters, and mapping atomistic thermodynamics onto CG potentials in a way that is both predictive and minimally overfit.

In this study, we introduce TEA (Temperature-dependent Energetics derived from hydration free energies), a physics-based framework that augment coarse-grained models with temperature-dependent potentials based on hydration free energies. Our approach includes three stages. First, we obtain the potential of mean forces between amino acid sidechain/backbone analogues from extensive all-atom MD simulations and quantify the temperature-dependent interactions. The accumulated simulation time for all-atom MD simulations is over 500 μs. Using these simulation data, we demonstrate that interaction strengths of the interacting species exhibit a linear a linear dependence on their hydration free energies. This linearity allows us to get the interaction strengths in the continuous temperature space. Moreover, we show that the heterotypic interactions among different species can be well approximated from the homotypic interactions using a combination rule, which enables scalable construction of a full interaction matrix. In the second stage, we leverage these physical insights and map the atomistic energetics to CG potentials via a linear relationship, which is governed by a single global scaling parameter, *γ*. This single free parameter allows us to integrate TEA into existing coarse-grained models with minimal reparameterization. Lastly, we fitted *γ* and validated this framework against independent experimental datasets. We demonstrate that the TEA-augmented CG potentials provide a transferable, physically interpretable route to simulate thermoresponsive IDPs with significant predictive power, successfully capturing LCST phase behavior and identifying sequence-specific outliers without relying on extensive parameter fitting to experimental data.

## Results

### Temperature-dependent homotypic interactions for amino acid sidechain analogues were obtained from all-atom MD simulations

To establish a thermodynamic basis for coarse-grained interactions, we quantified the temperature dependent interactions between amino acid sidechain/backbone analogues using extensive MD simulations. The model compounds used to mimic amino acid sidechain and backbone are listed in Table S1. The potential of mean forces (PMFs) for interacting pairs were obtained from umbrella sampling. We utilized the CHARMM36 force fields [19, 42-44] to get PMFs for homotypic pairs across a temperature range of 280 K to 380 K (Fig. S1). More details of the simulations can be found in Methods.

For hydrophobic analogues, such as Leu and Ile, the interaction strength increases with temperature, evidenced by a deepening minimum in the PMF curves (Fig. S1). Conversely, hydrophilic analogues, such as Ser and Thr, exhibit negligible temperature dependence (Fig. S1). For charged analogues, we isolated the solvent-mediated van der Waals (vdW) contributions by subtracting the direct coulombic interaction from the calculated PMF (details in Methods). The resultant PMFs after subtraction quantify the vdW interactions mediated by water molecules between the charged analogues. It is important to note that while we subtract the analytical Coulomb term, the direct electrostatic interactions still implicitly influence the resultant PMFs by affecting the water arrangement between the charged molecules. The goal of this decoupling is specifically to derive the coarse-grained force field parameters of the modified Lennard-Jones (LJ) potential of charged species, independent of the long-range electrostatics handled separately in the CG model. The resulting PMFs confirm the hydrophilic nature of charged analogues, showing minimal variation with temperature (Fig. S1).

To translate these PMFs into a metric suitable for CG parameterization, we defined the excess free energy (Δ*G*_*E*_) for an interacting pair:

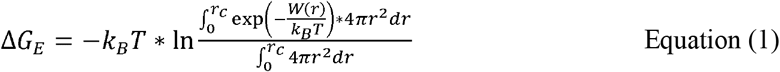

Where *W*(*r*) is the corresponding PMF, *r* is the inter-molecular distance, the cutoff *r*_*c*_ is set to 1.0 nm. Δ*G*_*E*_ provides a direct measure of the intrinsic interaction strength relative to a non-interacting ideal gas within the same volume, allowing for direct comparison across different chemical species. We note that the definition for Δ*G*_*E*_ is formally analogous to the standard binding free energy defined as 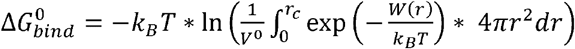,provided the region *r*< *r*_*c*_ is defined as the bound region. *V*^0^is the standard state volume 1.661 *nm*^3^, corresponding to a 1 M concentration. Consequently, Δ*G*_*E*_ and 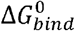 differ only by a molecule-independent value. Δ*G*_*E*_ is therefore ideal for calibrating the relative interaction energies required for fitting CG potential parameters.

Figure 1 shows Δ*G*_*E*_ as a function of temperatures for different homotypic interactions. As expected, Δ*G*_*E*_ decreases substantially for hydrophobic analogues (e.g., Ile and Phe) as temperature rises, indicating stronger attractions, whereas it shows no significant changes for hydrophilic residues (e.g., Ser and Gln). We further compared values of Δ*G*_*E*_ at 300 K for models for sidechain analogues with the corresponding hydrophobicity scales in several prominent CG models for simulating IDP phase separation. We found that Δ*G*_*E*_ (300 *K*) show high correlations with existing scales (Fig. S2). The correlation coefficients with HPS, HPS-Urry, HPS-FB, CALVADOS (M2), mpipi and mpipi-R are 0.63, 0.85, 0.65, 0.79, 0.64 and 0.62, respectively (Figure S2). We note that the HPS model assigns a *λ* value of zero to Arg, the key parameter indicating the van der Waals interaction strength with other residues. This assignment makes Arg a clear outlier in the Δ*G*_*E*_ (300 *K*) v.s. *λ* plot shown in Fig. S2. Without considering the charged species, Δ*G*_*E*_ (300 *K*) shows stronger correlations with these models, with coefficients of 0.78, 0.85, 0.70, 0.73, 0.69 and 0.66 for HPS, HPS-Urry, HPS-FB, CALVADOS (M2), mpipi and mpipi-R, respectively. Interestingly, correlations with the more physics-driven models like HPS-Urry and HPS are higher than those with more data-driven models including CALVADOS (M2), HPS-FB.

**Figure 1.**
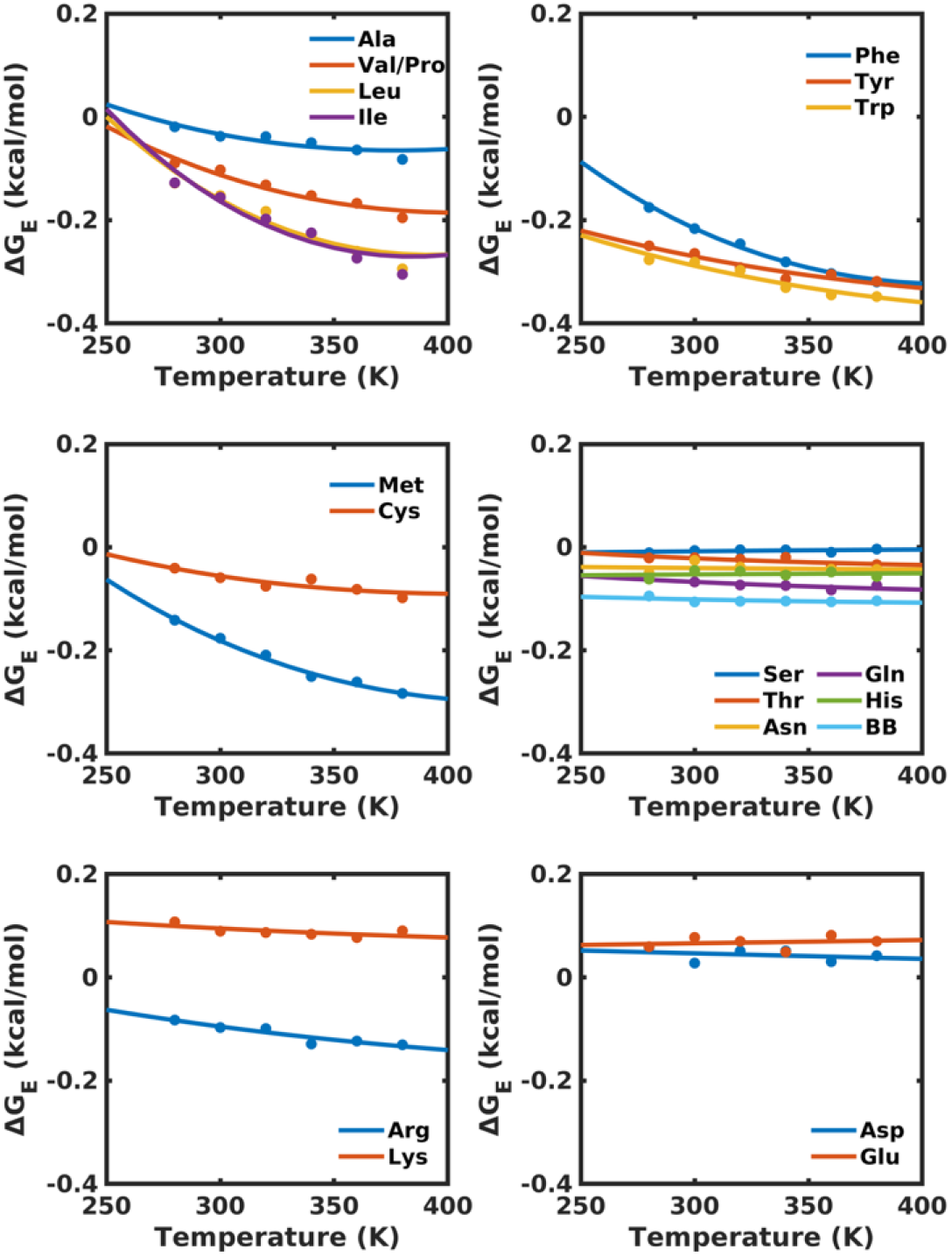
Temperature-dependent excess free energies Δ*G*_*E*_ for amino acid sidechain and backbone analogues. Solid spheres represent results calculated from potential of mean forces (PMFs), while solid lines indicate fits obtained by combining the integrated Gibbs-Helmholtz equation (Eq. 2) and with the linear scaling relationship (Eq. 3).

### Excess free energy Δ*G*_*E*_ can be derived from hydration free energyΔ*u*_*h*_

While umbrella sampling provides interaction data at discrete temperatures, a robust CG model requires continuous temperature dependence. All-atom models with implicit solvents, such as EEF1[45] and ABSINTH [24], account for temperature-dependent interactions by introducing a solvation term into the Hamiltonian. Inspired by this approach, we asked if Δ*G*_*E*_ can be approximated using hydration free energy.

Next, we calculated the hydration free energies Δ*u*_*h*_ for all analogous molecules at temperatures from 280 K to 380 K (Fig. 2). We fitted these data to the integrated version of the Gibbs-Helmholtz equation and obtained the temperature-dependent Δ*u*_*h*_:

**Figure 2.**
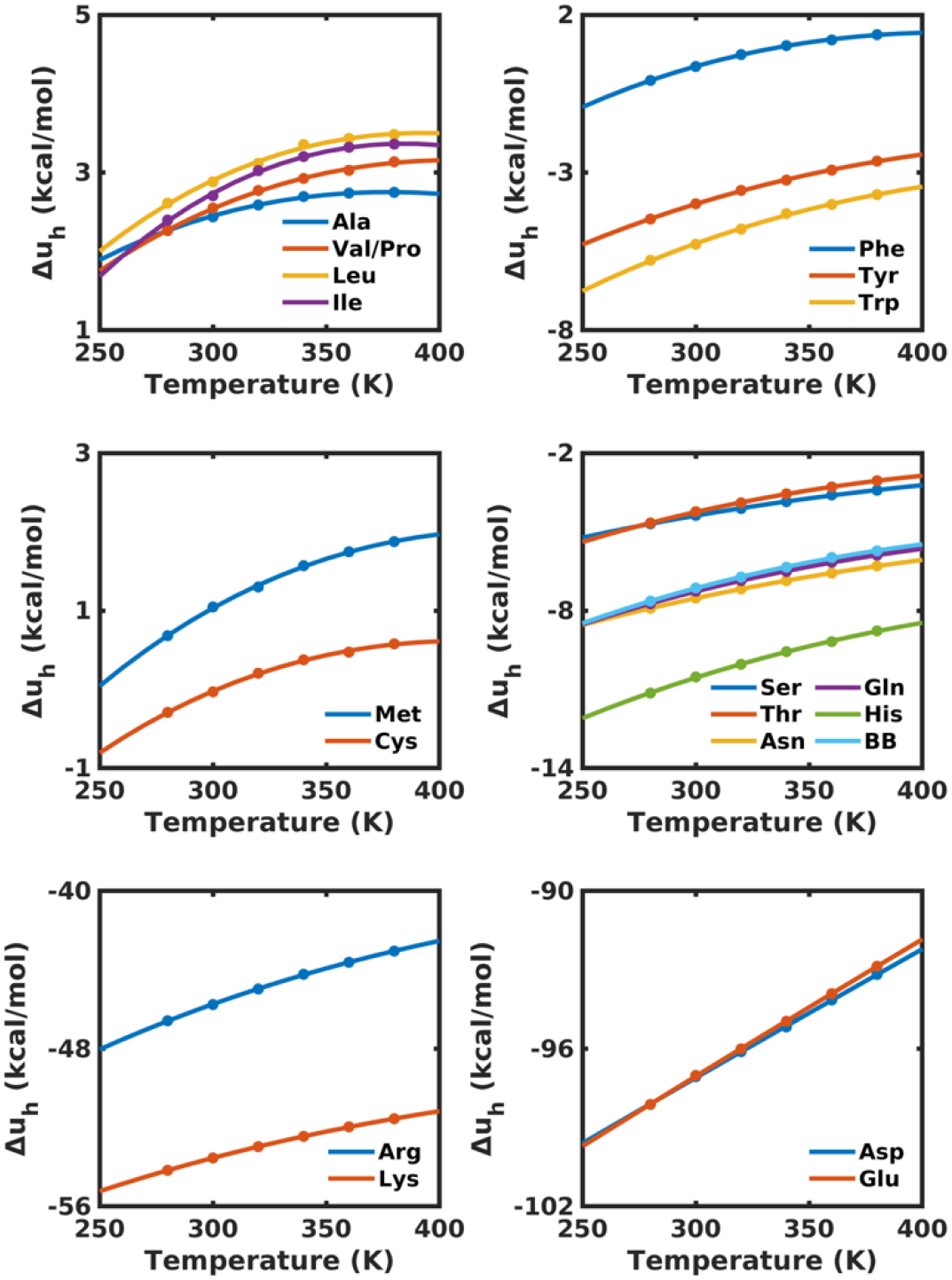
Temperature-dpendent hydration free energies Δ*µ*_*h*_ for amino acid sidechain and backbone analogues. Solid spheres represent results from free energy calculatoins in explicit solvent and solid lines are the fit to the integrated Gibbs-Helmholtz equation (Eq. 2).

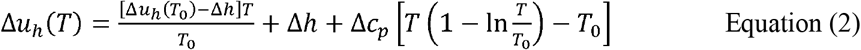

Here, Δ*h* is the hydration enthalpy at a reference temperature *T*_0_, Δ*c*_*p*_ is the change of molar heat capacity associated with the hydration process. *T*_0_ was set to 300 K. The fitted parameters for Δ*u*_*h*_ (*T*_0_), Δ*c*_*p*_ and Δ*h* are shown in Table S1. The fitted temperature-dependent Δ*u*_*h*_ (*T*) are shown by solid curves in Fig. 2.

A comparison of Δ*G*_*E*_ and Δ*u*_*h*_ reveals a striking linear correlation across all analogous molecules (Fig. 3A). This relationship allows us to express the interaction strength as a function of hydration free energies:

**Figure 3.**
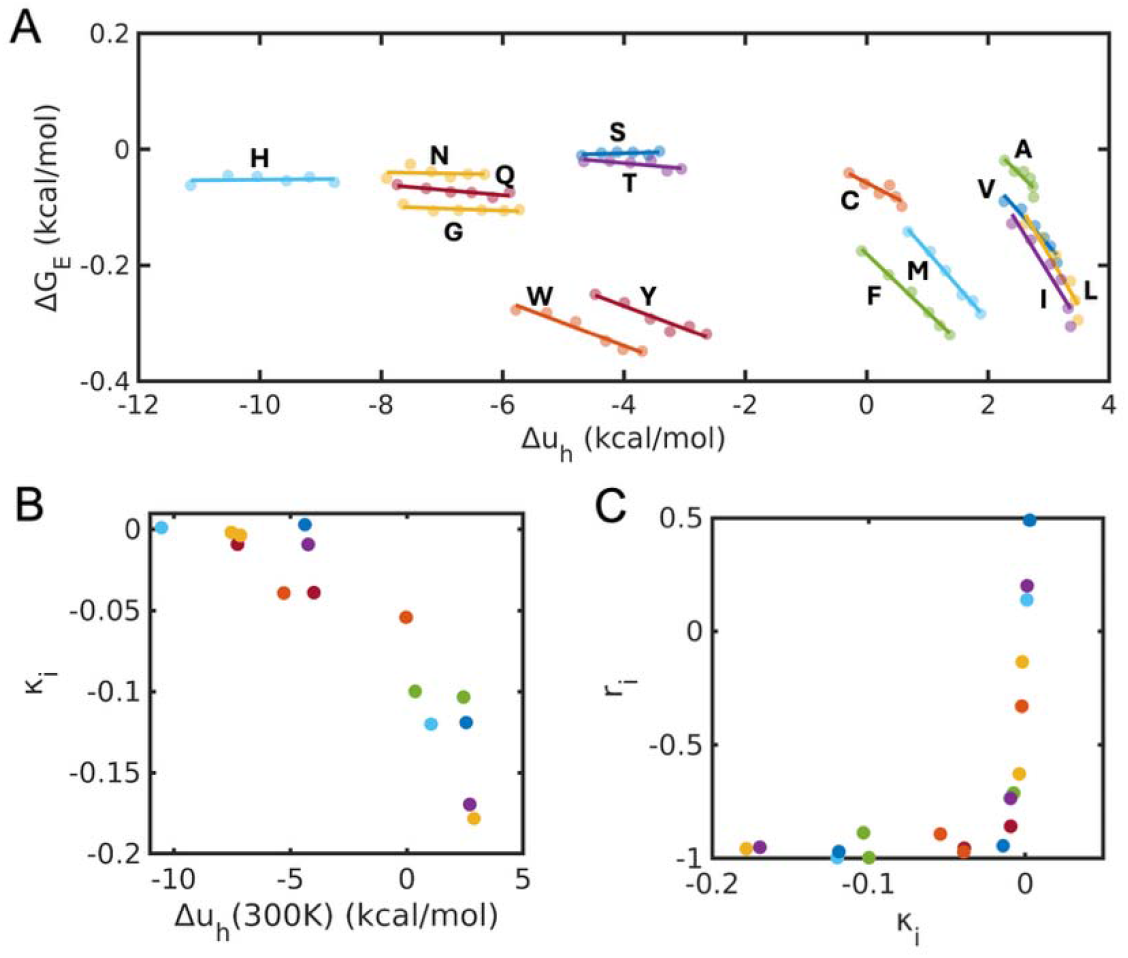
Excess free energies Δ*G*_*E*_ exhibit linear relationship with hydration free energies Δ*u*_*h*_. (A) Δ***G***_***E***_ plotted against Δ***u***_***h***_ for neutral amino acid side chain and backbone analogues. Solid spheres represent all-atom simulation data and solid lines are linear fits using Eq. 3. Data for charged species are provided in Fig. S3. (B) The residue-specific coefficients *κ*_*i*_ plotted as a function of hydration free energies at 300 K **Δ*u***_***h***_ 300*K*. (C) Dependence of the correlation coefficients *r*_*i*_ on the slope *κ*_*i*_.

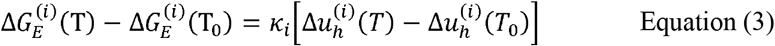

where *κ*_*i*_ is the residue-specific coefficient, quantifying the sensitivity of homotypic interactions to hydration changes. The mean absolute error of using Eq. 3 to fit the data is 0.007 kcal/mol. Values of *κ*_*i*_ and correlation coefficients *r*_*i*_ are listed in Table 1. Notably, for analogues with |*κ*_*i*_ | > 0.01, the corresponding |*r*_*i*_| values range from 0.88 to 0.98 (Fig. 3C), demonstrating a strong linearity between Δ*G*_*E*_ and Δ*u*_*h*_. However, it is important to note that the correlation coefficient is not always a reliable measure of linearity when the slope (*κ*_*i*_) is near zero. Specifically, for analogues with|*κ*_*i*_ | < 0.01, *r*_*i*_ spans from −0.86 to 0.50, while the mean absolute errors for these data points remain within 0.005 kcal/mol.

**Table 1.** Linear regression parameters correlating excess free energies (Δ*G*_*E*_ with hydration free energies (Δ*u*_*h*_). The slope *κ*_*i*_ and Pearson correlation coefficient *r*_*i*_ are provided for each residue analogue. They are derived from fitting Eq. 3 to all-atom simulation data. The mean absolute error indicates the deviation of the linear fit from the calculated values.

| Residue/unit | $\kappa_i$ | $r_i$ | Mean absolute error |
| --- | --- | --- | --- |
| Ala | -0.103 | -0.887 | 0.008 |
| Val/Pro | -0.119 | -0.970 | 0.008 |
| Leu | -0.178 | -0.957 | 0.015 |
| Ile | -0.170 | -0.951 | 0.017 |
| Met | -0.120 | -0.996 | 0.004 |
| Phe | -0.100 | -0.997 | 0.003 |
| Cys | -0.054 | -0.893 | 0.006 |
| Tyr | -0.039 | -0.955 | 0.006 |
| Trp | -0.039 | -0.971 | 0.006 |
| Ser | 0.003 | 0.489 | 0.002 |
| Thr | -0.009 | -0.735 | 0.004 |
| Asn | -0.002 | -0.134 | 0.006 |
| Gln | -0.009 | -0.859 | 0.003 |
| His | 0.001 | 0.139 | 0.006 |
| Backbone<br>/Gly | -0.004 | -0.628 | 0.002 |
| Arg | -0.014 | -0.944 | 0.005 |
| Arg<br>(charmm36) | 0.059 | 0.976 | 0.012 |
| Lys | -0.007 | -0.712 | 0.006 |
| Asp | -0.002 | -0.329 | 0.010 |
| Glu | 0.001 | 0.201 | 0.009 |

Figure 3A shows that hydrophobic analogues with relatively high Δ*u*_*h*_ values exhibit strongly negative *κ*_*i*_ values, confirming that their temperature-dependent attraction is driven by the entropic gain of dehydration. In contrast, hydrophilic analogues show *κ*_*i*_ values near 0, suggesting a weak dependence of interaction strength on hydration. Although there is a strong linear relationship between Δ*G*_*E*_ and Δ*u*_*h*_ for individual species, the relationship across different species is complex. Specifically, we cannot use a single universal κ value to connect Δ*G*_*E*_ and Δ*u*_*h*_ for all analogues.

Figure 3B shows the relationship between *κ*_*i*_ and Δ*u*_*h*_ (300 *K*) for non-charged analogues. Here, Δ*u*_*h*_ (300 *K*) reflects the hydrophobicity for each analogue. As the analogue become more hydrophilic, the dependence of Δ*G*_*E*_ on Δ*u*_*h*_ weakens, with *κ*_*i*_ approaching 0 when Δ*u*_*h*_ (300 *K*) falls below −6 kcal/mol. This suggests that in the highly hydrophilic regime, interactions become effectively temperature independent. Additionally, in this regime, the change in Δ*u*_*h*_ from 280 K to 380 K is comparable to or even larger than that in the hydrophobic regime, demonstrating a decoupling between Δ*G*_*E*_ and Δ*u*_*h*_. Furthermore, the data points in Fig. 3B appear to follow a master curve, suggesting the existence of a general physical principle governing the relationship between *κ*_*i*_ and Δ*u*_*h (300 K)*_.

By combining Eq. 2 and Eq. 3,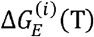 can be quantified within the continuous temperature space. Using the parameters for Δ*u*_*h*_ from all-atom simulations (Table S1) and the residue-specific *κ* values (Table 1), the strength of homotypic interactions can be directly obtained from hydration free energies.

### Heterotypic interactions are accurately predicted by a simple combination rule

Although homotypic interactions can be calculated based on hydration free energy, constructing the full interaction matrix for all residue pairs via umbrella sampling is computationally intractable. In all-atom MD simulations, combination rules are commonly used to approximate nonbonded heterotypic interactions between different atoms [46]. However, it remains unclear whether these simple rules hold validity at the molecular level. Many CG models, such as Martini 2 [47], HPS-FB [31] and CALVADOS [27], assume the validity of combination rules in their parameterization. Conversely, models like Martini 3 [48] and mpipi [28] employ pair-specific parameters for heterotypic interactions, allowing more parameterization flexibility. To address this uncertainty directly, we performed large-scale molecular simulations to evaluate whether heterotypic interactions among analogues can be approximated from homotypic data using standard combination rules.

We computed PMFs for 43 randomly selected heterotypic pairs at 300 K (Fig. S4-S8). These 43 pairs cover 5 distinct interaction types: hydrophilic-hydrophilic, hydrophobic and-hydrophilic, hydrophobic-hydrophobic, charged-charged, and cation-aromatic interactions (Table S3). We then compared the directly calculated excess free energy, 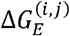,for a pair *I* and *j* against the arithmetic mean of their homotypic counterparts,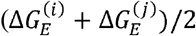.

As shown in Fig. 4, the direct calculations and the arithmetic means are highly correlated. This confirms that heterotypic interactions can be accurately approximated using a simple combination rule. Consequently, the full matrix of temperature-dependent interactions can be constructed solely from the hydration free energies of individual analogues, bypassing the need for exhaustive pairwise sampling.

**Figure 4.**
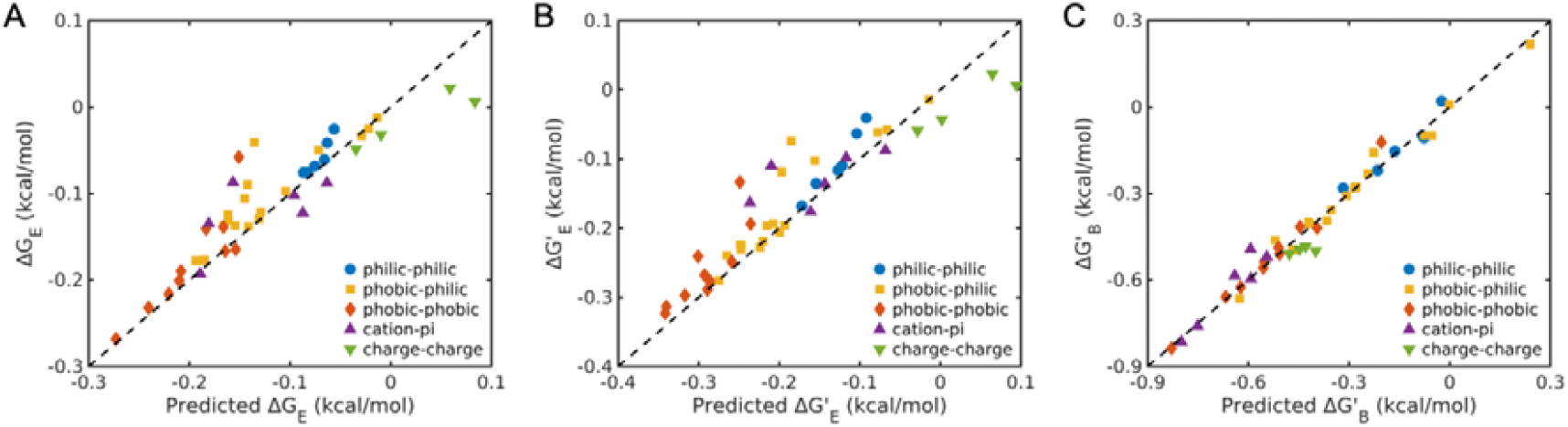
Validation of the combination rule for predicting heterotypic interactions. The plot compares values predicted using the combination rule (*x*-axis) against values calculated directly from all-atom simulations (*y*-axis) for **(A)** excess free energies Δ***G***_***E***_ calculated using a uniform cutoff 10 of Å, **(B)** excess free energies 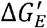 caculated using molecular-specific cutoffs, and **(C)** binding free energies 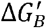 calculated using molecule-specific cutoffs. Molecule-specific cutoff values are listed in Table S2. Data points are colored by interaction type: hydrophilic-hydrophilic (blue), hydrophobic-hydrophilic (red), hydrophobic-hydrophobic (yellow), cation-*π* (purple), and charge-charge (green). The dashed line represents perfect agreement (*y = x*).

### Mapping temperature-dependent atomistic energetics to coarse-grained (CG) potentials

We next aim to integrate the temperature-dependent atomistic energetics to CG potentials. We chose the Hamiltonian form common to HPS [26] and HPS-based models [27, 31, 49].

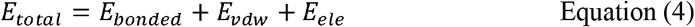

*E*_*vdw*_ is represented by the Ashbaugh-Hatch function:

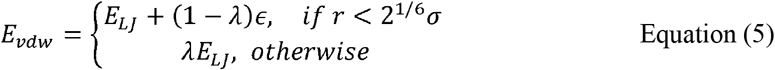

*E*_*LJ*_ is the standard Lennard-Jones potential. is 0.2 kcal/mol. The critical parameter in these models is the residue-specific stickiness parameter *λ*. A larger *λ* value represents stronger attractions. In the HPS and HPS-based models, *λ* values are constant across different temperatures. Temperature-dependent *λ* values were introduced using a parabolic function form in the HPS-T model [40].

To introduce temperature dependence, we mapped atomistic Δ*G*_*E*_ to the CG parameter *λ* via a linear transformation governed by a single global scaling factor *γ*:

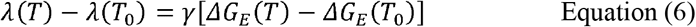

Here *T*_0_ is the reference temperature, 300 K. *γ* is a universal coefficient for all analogues. It accounts for the difference in energetics between all-atom model and coarse-grained model. This formulation allows us to augment existing HPS-based models, such as HPS-Urry [49], HPS-FB [31], and CALVADOS [27], with physics-based temperature dependence while retaining their original parameterization *λ* at the reference temperature *T*_0_. Our approach is similar to the recently developed mpipi-T model [41], which fitted linear coefficients between *λ* and Δ*u*_*h*_ using experimental data. Additionally, *γ* values for heterotypic interactions are fitted as well instead of using the combination rule. However, our method leveraged physical insights derived directly from extensive all-atom MD simulations to get the residue-specific *κ*_*i*_, leaving only a single free parameter *γ* to be fitted against experiment. This minimizes the risk of overfitting and ensures transferability. We term our approach TEA, abbreviated for “**T**emperature-dependent **E**nergetics derived from hydr**A**tion free energies”.

### TEA-augmented CG models robustly capture thermoresponsive phase behavior of IDPs

We next incorporated TEA to three CG models HPS-Urry [49], HPS-FB [31] and CALVADOS [27]. Then, we calibrated the scaling factor *γ* using experimental data on the thermoresponsive behaviors of resilin-like polypeptides (RLPs) and elastin-like polypeptides (ELPs) reported by Quiroz and Chilkoti [50]. RLPs and ELPs exhibit UCST- and LCST-type phase separation behaviors, respectively. Previous studies have demonstrated a strong coupling between single-chain conformations and the phase separation behaviors of IDPs [51-57]. Specifically, IDPs with UCST-type behavior expand at higher temperatures, whereas LCST-type IDPs collapse as temperature increases. Based on this relationship, we performed single-chain simulations for 23 RLP sequences and 16 ELP sequences across temperature from 250 K to 400 K.

We found that all the three TEA-augmented CG models, including CALVADOS, HPS-Urry, and HPS-FB, achieved a clear separation between RLP and ELP sequences when *γ* was set between 2 and 3 (Fig. 5 & S9-10). As shown in Fig 5, the radius of gyration (R_g_) for RLP sequences increase with temperature, while ELP sequences undergo compaction, consistent with their respective UCST- and LCST-type phase behavior. To have a quantitative assessment of the fitting quality, we compared the simulated single-chain R_g_ values at 350 K expected, of these sequences with their cloud point temperatures (*T*_*clound*_) observed in experiments. As we observed a positive correlation between the single-chain R_g_ values and (*T*_*clound*_) for the ELP sequences and a negative correlation for the RLP sequences (Fig. 5C-D & Fig. S9-10).

**Figure 5.**
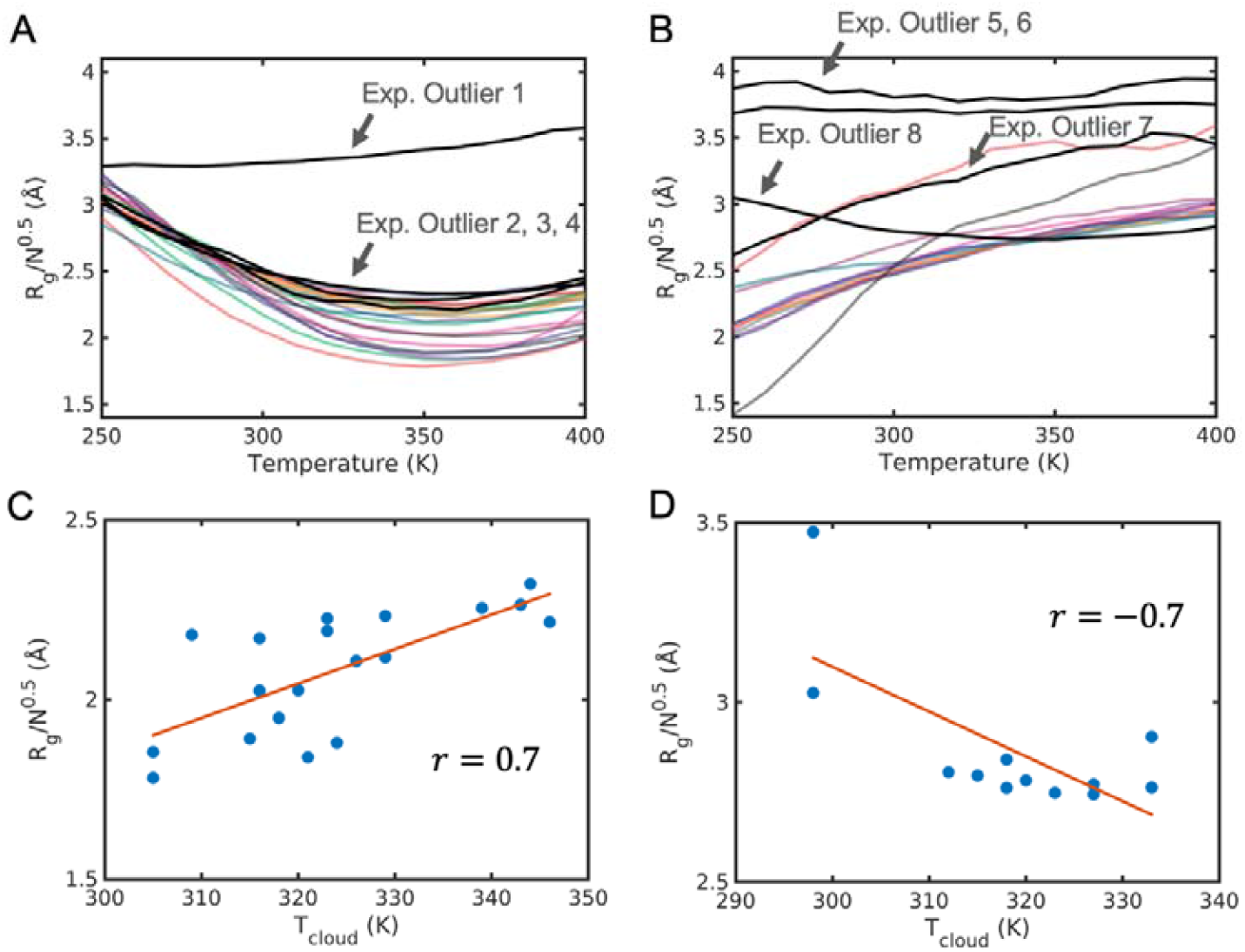
TEA-augmented HPS-Urry model distinguishes between LCST-type phase behavior in elastin-like polypeptides (ELPs) and UCST-type phase behavior in resilin-like polypeptides. Simulated single-chain R_g_ values at different temperatures for (A) ELPs and (B) RLPs. Experimental observed outliers are represented by black solid lines and indicated by arrows. Outliers 1-7 exhibit no phase separation within the temperature range from 20 ° C to 80 ° C in experiments. Outlier 8 is an RLP-like sequence, but undergoes LCST-type phase separation. The simulations correctly capture these trends. The normalized R_g_ values at 350 K for ELP sequences show a positive correlation with the experimentally measured cloud temperature T_cloud_ (C), while those for RLP sequences exibhit a negative

Remarkably, the TEA framework successfully identified outlier sequences reported in the experimental study by Quiroz and Chilkoti [50]. The RLP sequences (VRPVG)_25_, (TVPGAG)_55_, (VAPGVG)_20_, (APGVG)_45_, and ELP sequences (VPSDDYGVG)_29_, (VPSDDYGQG)_20_, (VPHSRNGG)_40_, (VPSTDYGVG)_12_, which are labeled as Exp. Outlier 1-7 in Fig. 5A-B, do not undergo phase separation within the experimental temperature range from 20 °C to 80 °C. Our simulations show that these sequences, especially Exp. Outlier 1, 5, 6, exhibit extended conformations with high R_g_ values at all the simulated temperatures from 250 K to 400 K, indicating the absence of phase separation. Importantly, our simulations show that the RLP sequence Exp. Outlier 8 exhibits LCST-type single-chain coil-to-globule transitions, which aligns its LCST-type phase behavior observed in experiments. Taken togather, these sequences were correctly identified by our approach. Their simulated coil-to-globule transition curves align with experimental observations, distinguishing them from the standard trends of their respective families.

Further validation was performed on five IDPs previously studied by Wuttke et al. including ProTa N, ProTa C, Integrase, Csp Tm, and *λ*-repressor [36]. It has been shown that the single-chain R_g_ values of these IDPs decreases with increasing temperature due to the temperature-dependent solvation. The TEA-augmented HPS-Urry and CALVADOS models successfully reproduced the temperature-dependent R_g_ values for Integrase and Csp Tm(Fig. 6A-B & Fig. S11-12). While the absolute R_g_ values for Integrase and Csp Tm from TEA-augmented HPS-FB model were overestimated, the degree to which they change with temperature matched the experimental trend (Fig. S13). For *λ*-repressor, all three models overestimated the R_g_ values, but result in the LCST type of single-chain coil-to-globule transition curves (Fig. 6C, S12-S13). Given that our current implementation of the TEA approach is specifically to fit the temperature dependence of *λ*values, we believe that its performance in capturing the temperature-dependent single-chain behavior is reasonable and acceptable. Further improvement can be achieved by adjusting *λ* values at the reference temperature in the original CG models, which is beyond the scope of this study. For the highly charged ProTa N and ProTa C, simulated Rg values from all three models shows negligible change with temperature (Fig. 6D-E, S11-S13). This inconsistency with experimental results is likely due to the absence of explicit ion effect in these implicit solvent CG models. The simulated Rg values at 350 K of these five sequences also show strong correlations with experimental results (Fig. 6F, S11-S13), further demonstrating the ability of the TEA approach in capturing the temperature-dependent single-chain behaviors. Taken together, these results demonstrate that the TEA approach can successfully captured the temperature-dependent single-chain behaviors of IDPs although it shows limitations on highly charged sequences due to the inherent limitations of the coarse-grained models.

**Figure 6.**
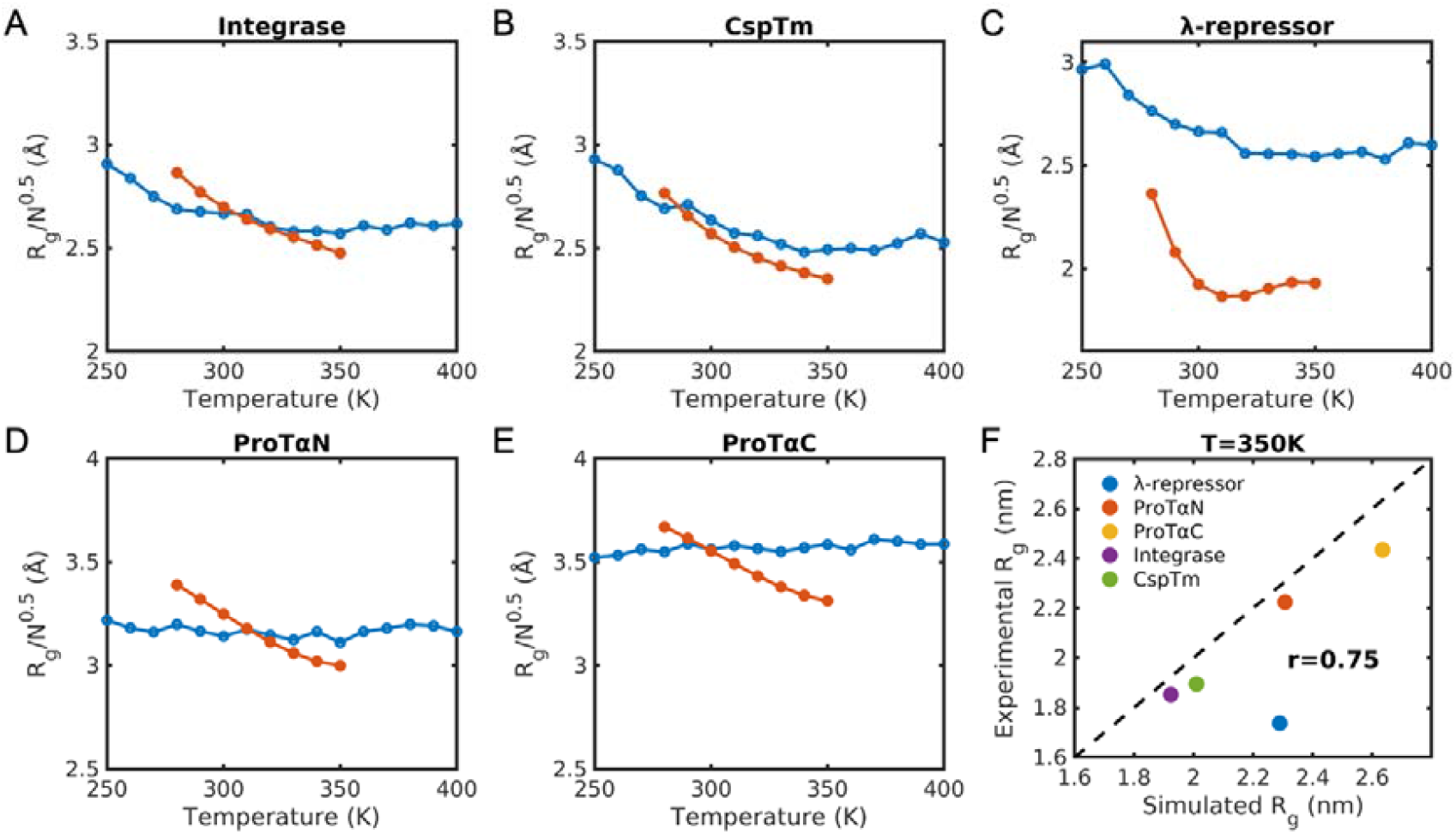
Comparison of simulated and experimental single-chain dimensions for five IDPs exihibiting LCST-type phase behavior. Temperature-dpendent R_g_ values from TEA-augmented HPS-Urry model with *γ*= 2 (blue curves) are compared with experimental measurements from Wuttke et al [36] (red curves) for (A) Integrase, (B) CspTm, (C) *λ*-repressor, (D) ProT *α* N, and (E) ProT *α* C. (F) Correlation between simulated and expeirmental R_g_ values at 350 K.

To validate the model’s ability to capture diverse phase behaviors, we computed phase diagrams for IDPs with varying thermoresponsive properties using the TEA-augmented HPS-Urry model. We selected three ELP and RLP sequences for analysis. As illustrated in Fig. 5A–B, the simulations correctly predict LCST-type phase separation for the ELP sequences and UCST-type behavior for the RLP sequences.

We extended this analysis to A1-LCD and its variants, which are known to exhibit UCST-type phase behavior [57, 58]. As shown in Figure 7C, the model successfully reproduces the experimental phase boundaries. Previous studies have established that tyrosine (Tyr/Y) acts as a stronger ‘sticker’ for phase separation than phenylalanine (Phe/F) [7, 58, 59]. Consequently, mutating Tyr to Phe increases the saturation concentration *C*_*sat*_. The TEA-augmented HPS-Urry model correctly captures the relative magnitudes of these shifts: the Tyr-enriched variant (−12F+12Y) exhibits a lower *C*_*sat*_, while the Phe-enriched variant (+7F-7Y) exhibits a higher *C*_*sat*_ compared to the wild type (WT). Although the relative trends are accurate, the absolute *C*_*sat*_ values differ from experimental data by approximately one order of magnitude, a limitation common to many coarse-grained models [60, 61]. Finally, consistent with experimental findings, the model demonstrates that sequences with higher aromatic content possess a stronger propensity for phase separation and lower saturation concentrations (Fig. 7D).

**Figure 7.**
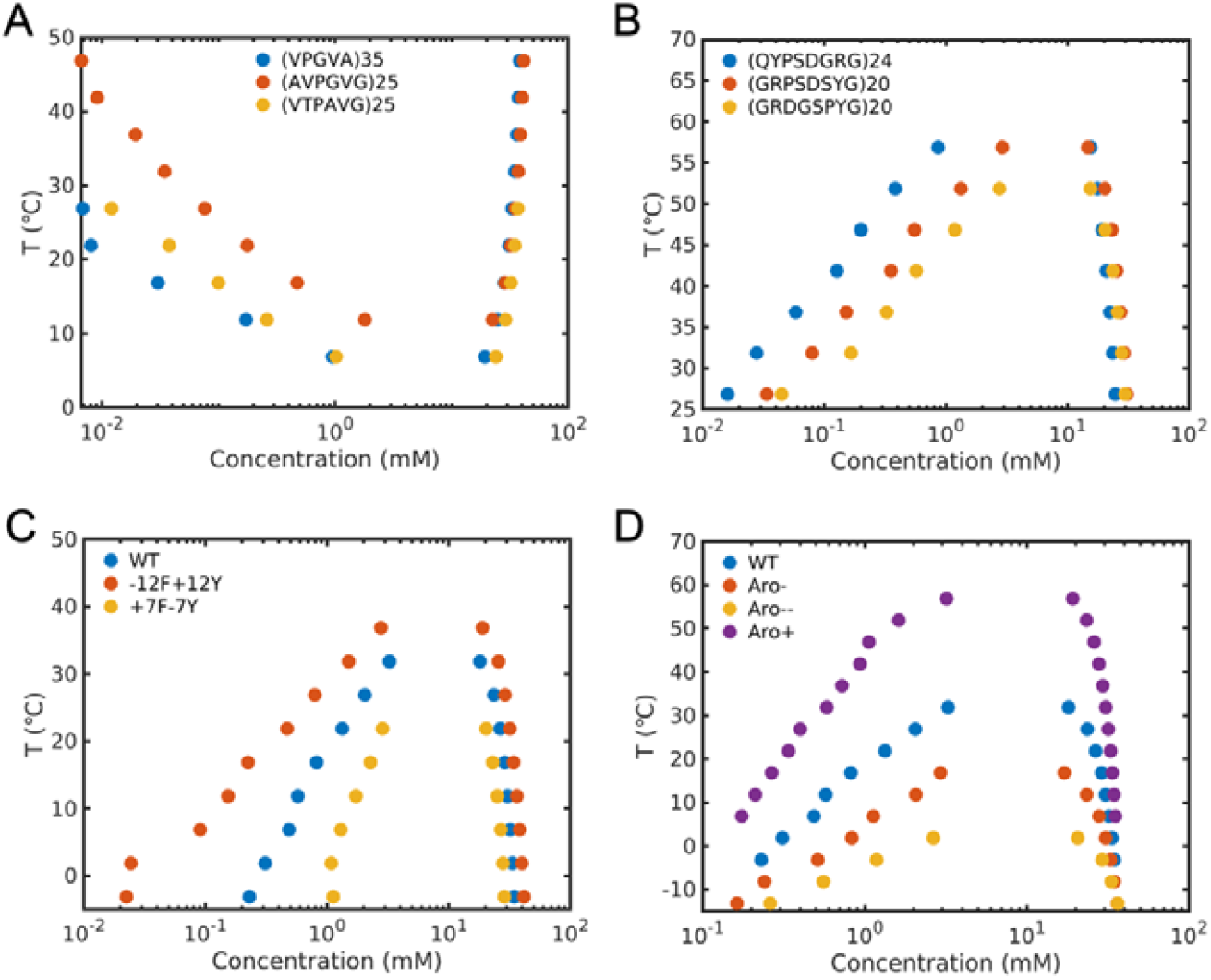
TEA augmented HPS-Urry model reproduces phase diagrams for diverse IDP sequences. Phase diagrams for (A) three ELP sequences, (B) three RLP sequences, (C) A1-LCD and two variants with altered number of Tyr and Phe, and (D) A1-LCD and three variants with different patterning of aromatic residues. The ELP and RLP sequences are selected from Quiroz and Chilkoti [50]. A1-LCD and its variants are from the work of Mittag’s group [57,58].

We further compared the phase diagrams obtained from TEA-augmented HPS-Urry with those from standard HPS-Urry with temperature-independent interactions for A1-LCD and its variants **(Fig. S14)**. Since the TEA-augmented HPS-Urry retains the original *λ* values at the reference temperature 300 K, the dilute and dense phase concentrations show minimal deviation around 300 K for all six sequences. The phase diagrams for the −12F+12Y variant are nearly identical between the two models, whereas the +7F-7Y variant exhibits a more significant deviation. This difference can be attributed to the stronger temperature dependence of Phe than Tyr as shown in Fig. 1. For the Aro- and Aro--variants, the dilute phase arms show mild variations while preserving the overall shape of the binodal. In contrast, the phase diagram for Aro+ displays a larger deviation and exabits a crossover at the reference temperature, which is likely driven by the strong cooperativity between aromatic residues due to their increased linear clustering. Overall, TEA-augmented HPS-Urry model preserves the ability to capture phase diagrams for UCST-type sequences while simultaneously enabling the description of LCST-type phase behavior.

## Discussion

In this study, we introduced the TEA framework to incorporate temperature dependent interactions to CG models for simulating thermoresponsive behaviors of IDPs. By employing extensive all-atom MD simulations, we successfully obtained the temperature-dependent energetics of amino acid sidechain analogues and further converted them to the coarse-grained potential. When augmented with TEA, coarse-grained models effectively distinguish between sequences that undergo LCST-type phase separation and UCST-type phase separation. Notably, this framework successfully identifies experimental outliers and capture both the LCST-type and UCST-type phase behaviors, demonstrating its significant predicative power.

### A physics-driven approach for transferability

Unlike purely data-driven approach that relies on fitting large experimental datasets, the TEA approach is fundamentally physics-based. It includes two stages: (a) quantify the temperature-dependent energetics from extensive all-atom MD simulations; (b) mapping this atomistic energetics to CG potentials via a single free parameter *γ*. This approach offers two advantages. First, this one-parameter fitting process minimizes the risk of overfitting when high-quality data is scare. Second, the physics-based approach enhances the model’s interpretability and transferability, allowing it to predict behaviors outside the training set while providing a clear understanding of its physical boundaries.

### All-atom energetics from extensive MD simulations

A key step of our approach is connecting the inter-residue interaction strength quantified by excess free energies Δ*G*_*E*_ with hydration free energies Δ*u*_*h*_. We establish a linear relationship between Δ*G*_*E*_ and Δ*u*_*h*_ by a residue-specific scaling factor *κ*_*i*_ as in Eq. 3. The linearity is well supported by the all-atom simulations (Fig. 3). Furthermore, we employed a combination rule to derive heterotypic interactions from homotypic data. The high correlation between simulated heterotypic interactions and those predicted by the combination rule confirm that the combination rule works well for these rigid sidechain analogues (Fig. 4).

To calculate Δ*G*_*E*_ from corresponding PMF curves, a uniform cutoff distance of 10 Å was used for all analogues. We note that within this range, the PMF curves generally exhibit a metastable state for most pairs indicated by a local minimum (Fig. S1), corresponding to the solvent mediated state without direct contact. This state is similar to the solvent-share ion pair (SIP) state observed in cation-anion interactions [62]. Applying a uniform cutoff yields a linear relationship between Δ*G*_*E*_ and the binding affinity 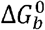 with a constant offset, confirming the robustness of this definition.

We subsequently evaluated a molecule-specific cutoff defined by the first energy barrier, which corresponds to the region of direct contact of the molecule pair. As shown in Figure S15, the resulting excess free energies 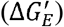 at 300 K correlates strongly with those derived using the 10 Å cutoff. The 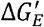 versus Δ*u*_*h*_ plot is shown in Fig. S16, with fitted parameters listed in Table S2. Notably, the residue-specific scaling factor 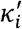 connecting 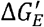 and Δ*u*_*h*_ are similar with the those using a uniform cutoff (Fig. S16D). Furthermore, the 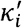 versus Δ*u*_*h*_ (300 *K*) plot in Fig. S16B follows a similar mater curve shown in Fig. 3B and the combination rule holds validity as well as shown in Fig 4B.

Surprisingly, when the residue specific cutoff distance is used to calculate the binding free energy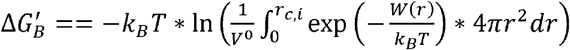, where *r*_*c,i*_ considers the direct contact, the combination rule exhibit exceptionally high fidelity (Fig. 4C). While the physical origin of this improved correlation warrants further investigation, the robustness of the results suggests our energetic derivation is not an artifact of the cutoff choice.

### Fit lambda values from all-atom energetics

We directly set the *λ* values at the reference temperature, *λ (T*_*0*_), to be the original *λ* values in the HPS-based models and obtain the temperature dependence of *λ* values from Δ*G*_*E*_ using a linear relationship (Eq. 6). By combining Eq. 2, 3 and 6, *λ* becomes an analytical function of thermodynamic observables Δ*u*_*h*_ (*T*_0_), Δ*c*_*p*_, Δ*h*., the scaling factor *κ*, and the fitting parameter *γ*. While the thermodynamic terms and *κ* are obtained directly from the all-atom simulation, *γ* is the sole tunable parameter obtained to match experimental data. This approach simplifies the fitting process significantly. We note that *λ* (*T*_0_) can be derived from Δ*G*_*E*_ (*T*_0_) in principle, which will introduce additional fitting parameters besides *γ*. Therefore, we leverage existing CG models to constrain the reference values, thereby maintaining a simple but efficient model with minimal free parameters.

We point out that the influence of coarse-grained bead radius *σ* on the interaction strength is ignored in parameterizing *λ*. A more rigorous alternative is to fit the *λ* values by connecting the all-atom Δ*G*_*E*_ with coarse-grained one 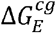 by a linear relationship. However, because 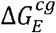 is not an elementary function of *λ* and depends on the residue-specific bead radius, such approach makes the fitting process not straightforward and requires numerical solutions. Our simplified method maintains tractability while effectively capturing the essential physics. It is important to note that while we fit the temperature-dependent potential for the one-bead-per-residue models, the underlying energetics are derived from sidechain analogues. Consequently, this implementation assumes that the backbone contribution is uniform across residues and that backbone-sidechain cooperativity is negligible. Future refinements could employ two-bead-per-residue models to explicitly decouple backbone and sidechain energetics.

Regarding the force field dependence, our current *κ*_*i*_ and Δ*u*_*h*_ values are derived using the Charmm forcefields. We believe that *κ*_*i*_ is an intrinsic thermodynamic quantity and therefore the choice of forcefields will not significantly influence the *κ*_*i*_ values, or alter the relative ranking of *κ*_*i*_ values. Moreover, experimental results of temperature-dependent hydration free energies are available for these 19 sidechain analogues [63-65], and temperature-dependent second virial coefficients can be measured in experiments to calibrate *λ* values [52, 66, 67]. Future iterations of TEA could incorporate this experimental data directly, which will render the framework force-field independent, potentially increasing its reliability.

### Robustness and limitations

The TEA framework is highly versatile and compatible with existing one-bead-per-residue models. In this study, we successfully incorporated the temperature-dependent energetics to the HPS-Urry, HPS-FB, and CALVADOS coarse-grained models. Although the single-chain R_g_ values at 300 K differ for these models due to their intrinsic parameterization, the trend of the temperature-dependent single-chain behavior remains consistent across these three models. This suggests that the derived temperature-dependent energetics are robust and transferable. We anticipate that the TEA framework can be extended to other coarse-grained models for modeling biomolecular phase separation, such as mpipi [28], AWSEM-IDP [68], COCOMO [69], MOFF [70], and HyRes [71].

However, we do not expect that a coarse-grained model can capture the thermoresponsive behavior of all types of IDPs if the necessary molecular features are ignored in the design of the coarse-grained model. Indeed, we found that the current TEA implementation does not accurately model highly charged IDPs where ion effects are crucial. This is an expected consequence of using implicit solvent models lacking explicit ions. It is widely recognized that ion binding and releasing are important driving forces for polymer phase separation [72-75], and recent studies have demonstrated the necessity of explicit ions for simulating phase separation of highly charged biomolecules [76-78]. Therefore, extending the TEA approach to include explicit ion representations will be a crucial step toward a comprehensive model of IDP phase behavior.

## Methods

### Details of all-atom molecular dynamics simulations

All simulations were performed using Gromacs 2024.2 package [79]. The CHARMM General Force Fields (CGenFF) [42-44] were used for model compounds, with the exception of the arginine analogue. We compared the Lennard-Jones parameters (*σ* and *ε*) and partial charges against those from CHARMM36m protein force field [19] (Fig. S15-S18). While values from these two force fields are nearly identical for most analogues, we observed a significant difference in the partial charges of guanidinium nitrogen in arginine analogue. Consequently, we adopted CHARMM36m parameters for arginine analogue to ensure consistency with protein parameterization. CHARMM modified TIP3P water model was used [80]. All simulations were first energy minimized using the steepest descent algorithm. This was followed by an equilibration phase of 0.5 ns in the NPT ensemble. Production simulations were subsequently performed in the NPT ensemble. Temperature was kept at the target value using v-rescale thermostat [81] with a coupling time constant of 0.1 ps for the umbrella sampling, while Langevin thermostat with the inverse friction constant of 2 ps was used for the hydration free energy calculations. Pressure was maintained at 1 bar by the Parrinello-Rahman barostat [82, 83] with a coupling constant of 2 ps. The distance cutoff for both short-range van der Waals and electrostatic interactions were 1.2 nm. Periodic boundary condition was applied and the Particle Mesh Ewald (PME) method [84, 85] was used to calculate the long-range electrostatic interactions. The integration time step is 2 fs. The integrators for umbrella sampling and hydration free energy calculations are *md* and *sd*, respectively.

### Umbrella sampling for getting potential of mean forces (PMFs)

Umbrella sampling [86] combined with the weighted histogram analysis method (WHAM) [87, 88] were used to obtain PMFs. For each system, nine windows with centers from 0.4 nm to 2.0 nm with an interval of 0.2 nm were used. In each window, 4 independent simulations were performed, each with 100 ns long. Therefore, the accumulated simulation time for the 19 homotypic interaction pairs at 6 different temperatures is 410.4 μs, and the accumulated simulation time for the 43 heterotypic interaction pairs at 300 K is 154.8 μs. The total simulation time for PMF calculations is 565.2 μs. The harmonic restraint potential with a force constant of 1.0 *kcal* · *mol*^−1^ · Å^−2^ was applied to the distance between key functional groups in the model compounds. Key functional groups or atoms for each residue is listed in Table S4.

### Hydration free energy calculations

Slow-growth methods were used for simulations and the Bennett Acceptance Ratio (BAR) method [73] implemented in GROMACS package was used to calculate the free energy difference between two adjacent *λ* windows. *λ* values for the van der Waals interactions and electrostatic interactions for each window are[*λ*_*vdw*_, *λ*_*ele*_]=[0, 0], [0.1, 0], [0.2, 0], [0.3, 0], [0.4, 0], [0.5, 0], [0.6, 0], [0.7, 0], [0.8, 0], [0.9, 0], [1.0, 0], [1.0, 0.1], [1.0, 0.2], [1.0, 0.3],[1.0, 0.4], [1.0, 0.5], [1.0, 0.6], [1.0, 0.7], [1.0, 0.8], [1.0, 0.9], [1.0,1.0]. For each *λ* window, 10 ns simulations in the NPT ensemble were performed.

### Coarse-grained molecular dynamics simulations

The single-chain simulations were performed using the LAMMPS 2-Aug-2023 simulation package [89]. For each sequence, a 2-μs long simulation for sequences with less than 200 residues and a 4μs-long simulation for longer sequences were performed to calculate the R_g_ values at each target temperature. Initial 50 ns was discarded as equilibration. Langevin integrator with a timestep of 10 fs and friction coefficient of 0.5 ps^−1^ was used for the simulations. All interactions were truncated at 4 nm. The simulation box is 80 × 80 × 80*nm*^3^. Temperature-dependent dielectric constants were used as in the CALVADOS [27] and mpipi model [28].

The multiple-chain simulations for calculating phase diagrams were performed using the CALVADOS package [90]. The slab simulation box with the size of 15 × 15 × 150 *nm*^3^ was used. 100 proteins were inserted into the box with the overall concentration of 4.92 mM. For each temperature, 10 μs production simulations were performed. The first 1 μs was treated as equilibration and the rest 9 μs was used to calculate the dilute and dense phase concentrations. The default values in CALVADOS package [90] were used for all other parameters.

## Author Contributions

X.Z. conceived the idea, designed the study, analyzed simulation results, wrote the manuscript. C.F. performed the simulations, analyzed simulation results, and revised the manuscript.

## Acknowledgements

This work is supported by the funding from Research Grant Council of Hong Kong (Grant No. 22302823), National Natural Science Foundation of China (Grant No. 22303073), Hong Kong Baptist University (RC-FNRA-IG/22-23/SCI/03). We thank Dr. Yong Wang from Zhejiang University for the helpful discussions. All simulations were performed on the MGPU Cluster at the High-Performance Cluster Computing Centre, Hong Kong Baptist University.

