## Supporting Information for "Simulating Thermoresponsive Behavior of Disordered Proteins with Temperature-Dependent Coarse-Grained Potentials Derived from Hydration Free Energies"

for

**Table S1. Thermodynamic parameters for hydration free energies.** Parameters are obtained by fitting the temperature-dependent hydration free energies for the 19 amino acid sidechain/backbone analogues to the integrated version of the Gibbs-Helmholtz equation.

| Residue/unit | Model compound | $\Delta u_h(300K)$ kcal/mol | Δh kcal/mol | $\Delta c_p$ cal/molK |
|---|---|---|---|---|
| Ala | Methane | 2.45 | 0.02 | 35.30 |
| Val/Pro | Propane | 2.54 | -1.13 | 40.50 |
| Leu | 2-methylpropane | 2.90 | -1.17 | 50.49 |
| Ile | *n*-butane | 2.74 | -1.93 | 62.68 |
| Met | Ethyl methyl thioether | 1.03 | -3.75 | 43.66 |
| Phe | Toluene | 0.37 | -5.71 | 63.67 |
| Cys | Methanethiol | -0.02 | -3.69 | 39.10 |
| Tyr | *p*-Cresol | -4.00 | -10.60 | 41.91 |
| Trp | 3-methylindole | -5.25 | -12.94 | 50.57 |
| Ser | Methanol | -4.38 | -8.83 | 21.47 |
| Thr | Ethanol | -4.24 | -10.10 | 37.74 |
| Asn | Acetamide | -7.53 | -13.01 | 24.54 |
| Gln | Propionamide | -7.27 | -13.80 | 36.41 |
| His | 4-methylimidazole | -10.55 | -18.72 | 41.86 |
| Backbone/Gly | *N*-methylacetamide | -7.15 | -14.06 | 41.81 |
| Arg | *n*-propylguanidine | -45.74 | -57.99 | 58.02 |
| Lys | 1-butylamine | -53.54 | -62.55 | 41.93 |
| Asp | Acetic acid | -97.11 | -111.95 | 3.49 |
| Glu | Propionic acid | -97.05 | -112.90 | 4.68 |

**Table S2. Linear regression parameters between $\Delta G'_E$ and $\Delta u_h$.** $\Delta G'_E$ is the excess free energy calculated using the molecular-specific cutoff $r_{cut,i}$. $\kappa'_i$ is the slope and $r'_i$ is the correlation coefficient for analogue *i*.

| **Residue/unit** | $\kappa'_i$ | $r'_i$ | **Mean absolute error (kcal/mol)** | **Cutoff $r_{cut,i}$ (Å)** |
|---|---|---|---|---|
| Ala | -0.392 | -0.943 | 0.019 | 6.0 |
| Val/Pro | -0.260 | -0.981 | 0.013 | 7.4 |
| Leu | -0.333 | -0.974 | 0.022 | 7.6 |
| Ile | -0.264 | -0.962 | 0.024 | 8.2 |
| Met | -0.138 | -0.997 | 0.005 | 9.6 |
| Phe | -0.144 | -0.996 | 0.005 | 8.6 |
| Cys | -0.177 | -0.974 | 0.009 | 6.4 |
| Tyr | -0.055 | -0.969 | 0.008 | 8.8 |
| Trp | -0.031 | -0.962 | 0.006 | 11.4 |
| Ser | -0.002 | -0.886 | 0.000 | 6.6 |
| Thr | -0.013 | -0.776 | 0.005 | 9.0 |
| Asn | -0.012 | -0.497 | 0.009 | 6.8 |
| Gln | -0.021 | -0.941 | 0.004 | 7.6 |
| His | -0.002 | -0.299 | 0.005 | 7.4 |
| Backbone/Gly | -0.014 | -0.862 | 0.005 | 7.8 |
| Arg | -0.014 | -0.940 | 0.005 | 9.8 |
| Arg (charmm36m) | 0.063 | 0.979 | 0.012 | 9.8 |
| Lys | -0.008 | -0.761 | 0.006 | 10.3 |
| Asp | -0.001 | -0.122 | 0.010 | 9.2 |
| Glu | 0.002 | 0.354 | 0.009 | 9.2 |

**Table S3. Categorization of 43 heterotypic interaction pairs.** PMFs at 300 K for these pairs are calculated from all-atom molecular simulations. The one-letter amino acid code was used to represent the corresponding analogue.

| **Type** | **Heterotypic pairs** |
|---|---|
| Hydrophilic-Hydrophilic | G-N; G-Q; G-S; G-T; G-H; G-C |
| Hydrophobic-Hydrophilic | S-T; Y-N; Y-Q; Y-S; Y-T; G-A; G-M; G-Y; G-V; G-L; G-I; S-A; T-A; G-W; G-F; Y-H; Y-C |
| Hydrophobic-Hydrophobic | Y-A; Y-M; Y-V; Y-L; Y-I; M-L; M-I; L-I; Y-W; Y-F |
| Cation-$\pi$ | R-Y; R-W; R-F; K-Y; K-W; K-F |
| Charge-Charge | R-D; R-E; K-D; K-E |

**Table S4. Key functional groups in each analogue used to calculate the inter-molecular distance for applying the harmonic potential in umbrella sampling.**

| Residue/unit | Model compound | Functional group or atom |
|---|---|---|
| Ala | Methane | Carbon atom |
| Val/Pro | Propane | Carbon atom in the middle |
| Leu | 2-methylpropane | Carbon atom at the center |
| Ile | *n*-butane | Carbon atom in the middle |
| Met | Ethyl methyl thioether | Sulfur atom |
| Phe | Toluene | Aromatic ring |
| Cys | Methanethiol | Sulfur atom |
| Tyr | *p*-Cresol | Aromatic ring |
| Trp | 3-methylindole | Aromatic ring |
| Ser | Methanol | Oxygen atom |
| Thr | Ethanol | Oxygen atom |
| Asn | Acetamide | Carbon atom bonded to oxygen |
| Gln | Propionamide | Carbon atom two bonds away from oxygen |
| His | 4-methylimidazole | 5-membered ring |
| Backbone/Gly | *N*-methylacetamide | Carbon atom bonded to oxygen |
| Arg | *n*-propylguanidine | Carbon atom in guanidinium group |
| Lys | 1-butylamine | Nitrogen atom |
| Asp | Acetic acid | Carbon atom bonded to oxygen |
| Glu | Propionic acid | Carbon atom bonded to oxygen |

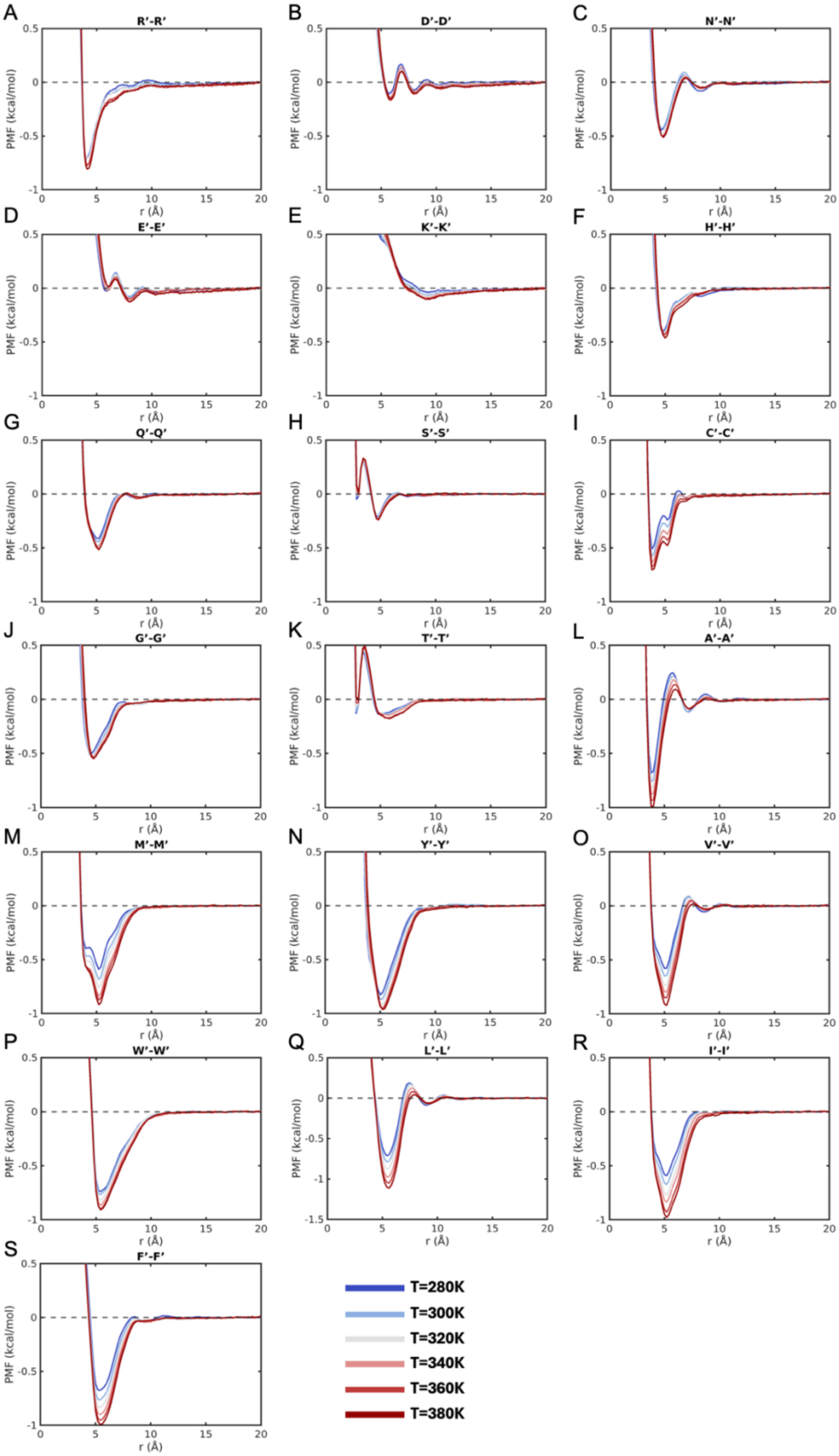


**Figure S1. Temperature-dependent potential of mean forces for the homotypic interactions of the 19 analogues.**

A

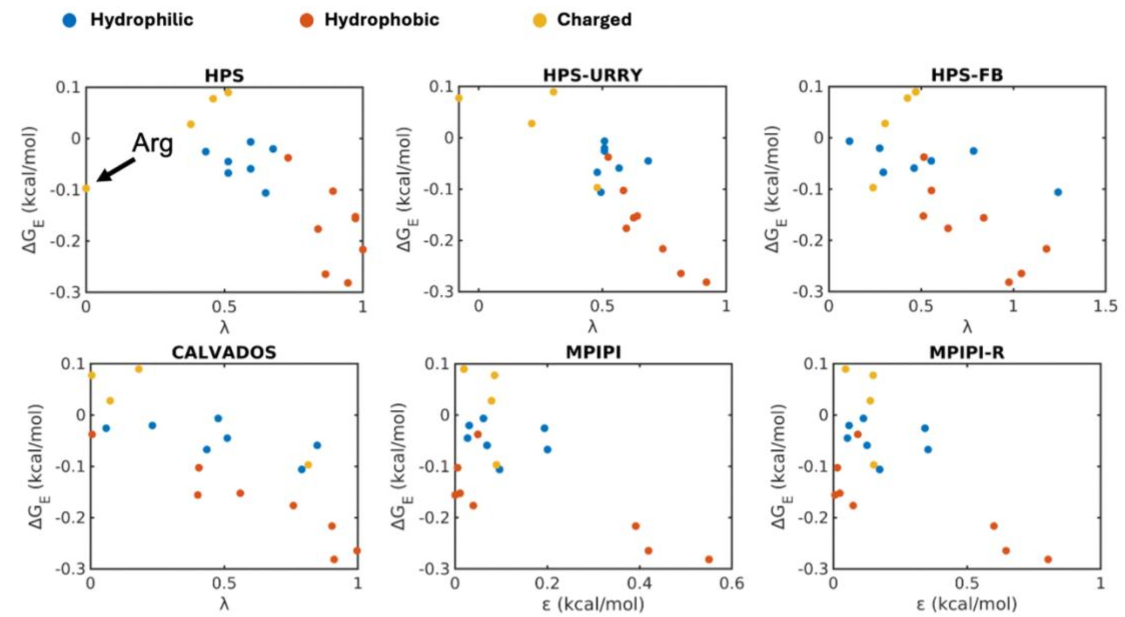


B

| | HPS | HPS-URRY | HPS-FB | CALVADOS | MPIPI | MPIPI-R |
|---|---|---|---|---|---|---|
| **All analogues** | 0.634 | 0.849 | 0.654 | 0.792 | 0.638 | 0.615 |
| **Without Charged** | 0.777 | 0.850 | 0.690 | 0.733 | 0.689 | 0.659 |

C

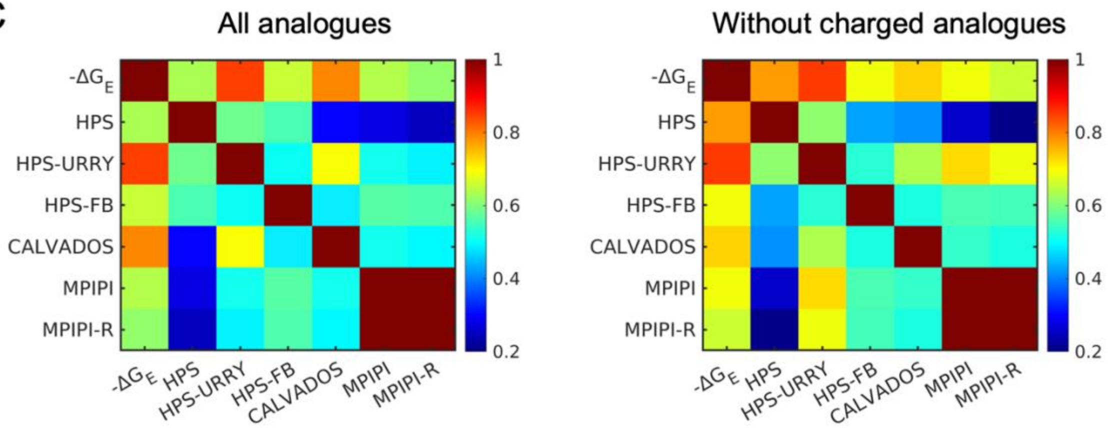


**Figure S2. Excess free energies at 300 K calculated from all-atom simulations, $\Delta G_E(300K)$, exhibit strong correlation with hydrophobicity scales in exsiting CG models.** (A) Scatter plot comparing $\Delta G_E(300K)$ against hydrophobicity scales in HPS, HPS-Urry, HPS-FB, CALVADOS, mpipi, and mpipi-R CG models, and (B) the corresponding correlation coefficients. (C) The cross correlation coefficient matrix across $\Delta G_E(300K)$ and hydrophobicity scales in these CG models.

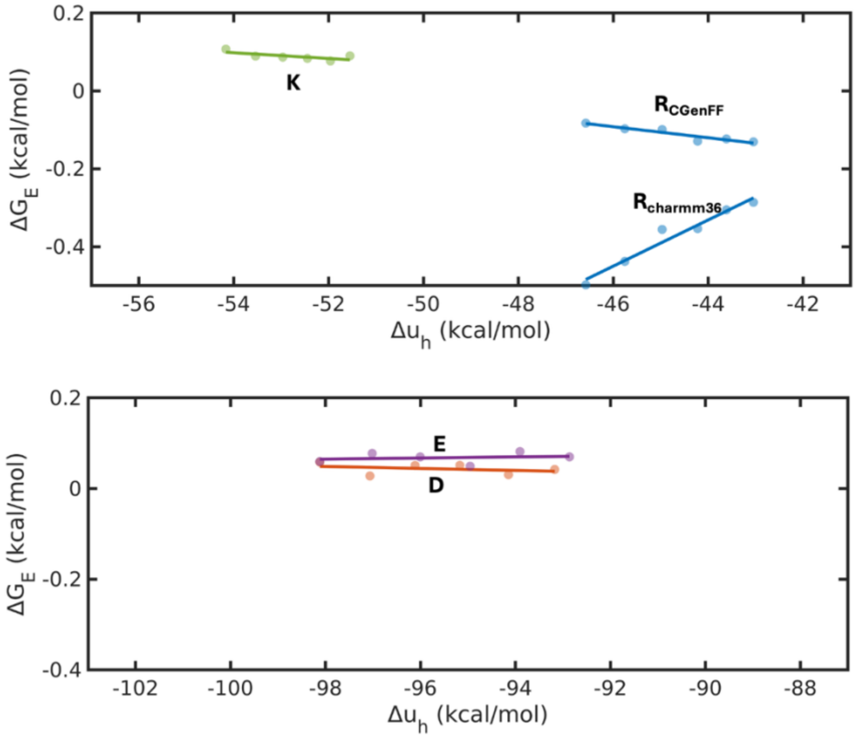


**Figure S3. $\Delta G_E$ versus $\Delta u_h$ for charged analogues.** Data for K, E, D, $R_{CGenFF}$ are from all-atom simulations using CGenFF, while those for $R_{charmm36}$ are from Charmm36m protein force field.

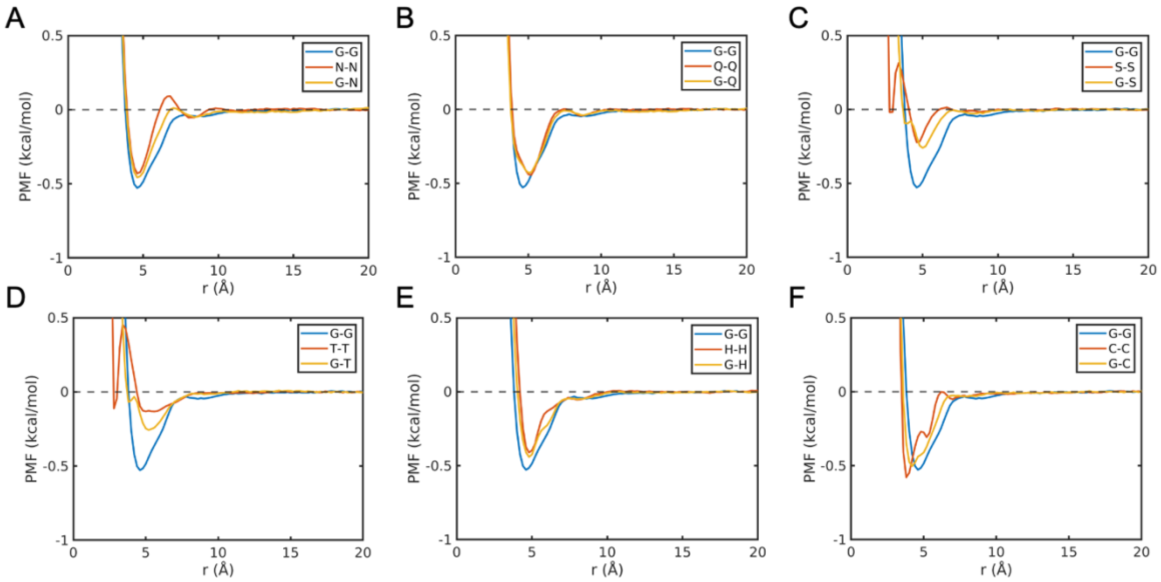


**Figure S4. PMFs for the 'hydrophilic-hydrophilic' heterotypic interactions at 300 K.** Heterotypic interactions are shown by yellow curves. For comparison, the corresponding homotypic interactions are shown in blue and red.

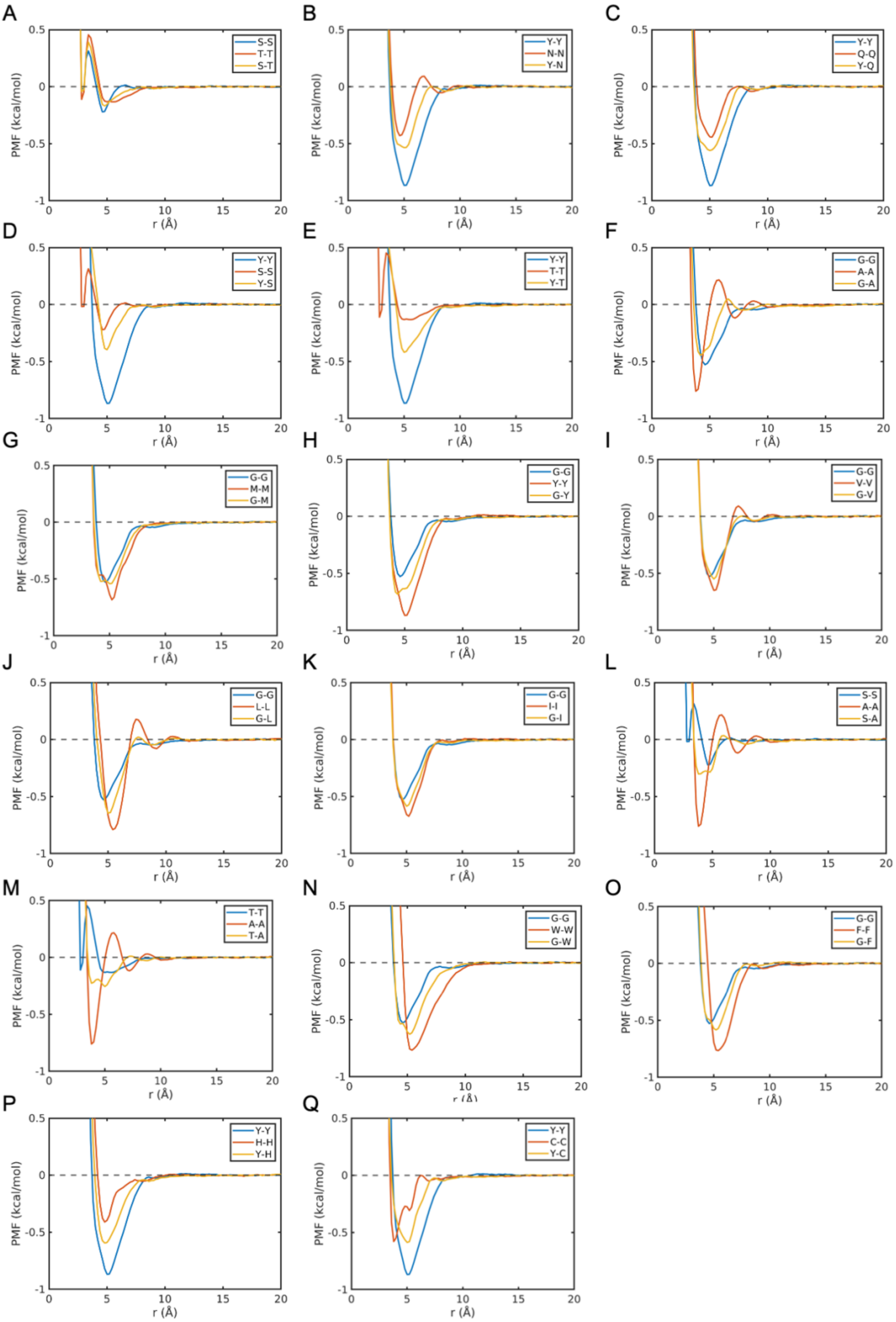


**Figure S5. PMFs for the 'hydrophobic-hydrophilic' heterotypic interactions at 300 K.** Heterotypic interactions are shown by yellow curves. For comparison, the corresponding homotypic interactions are shown in blue and red.

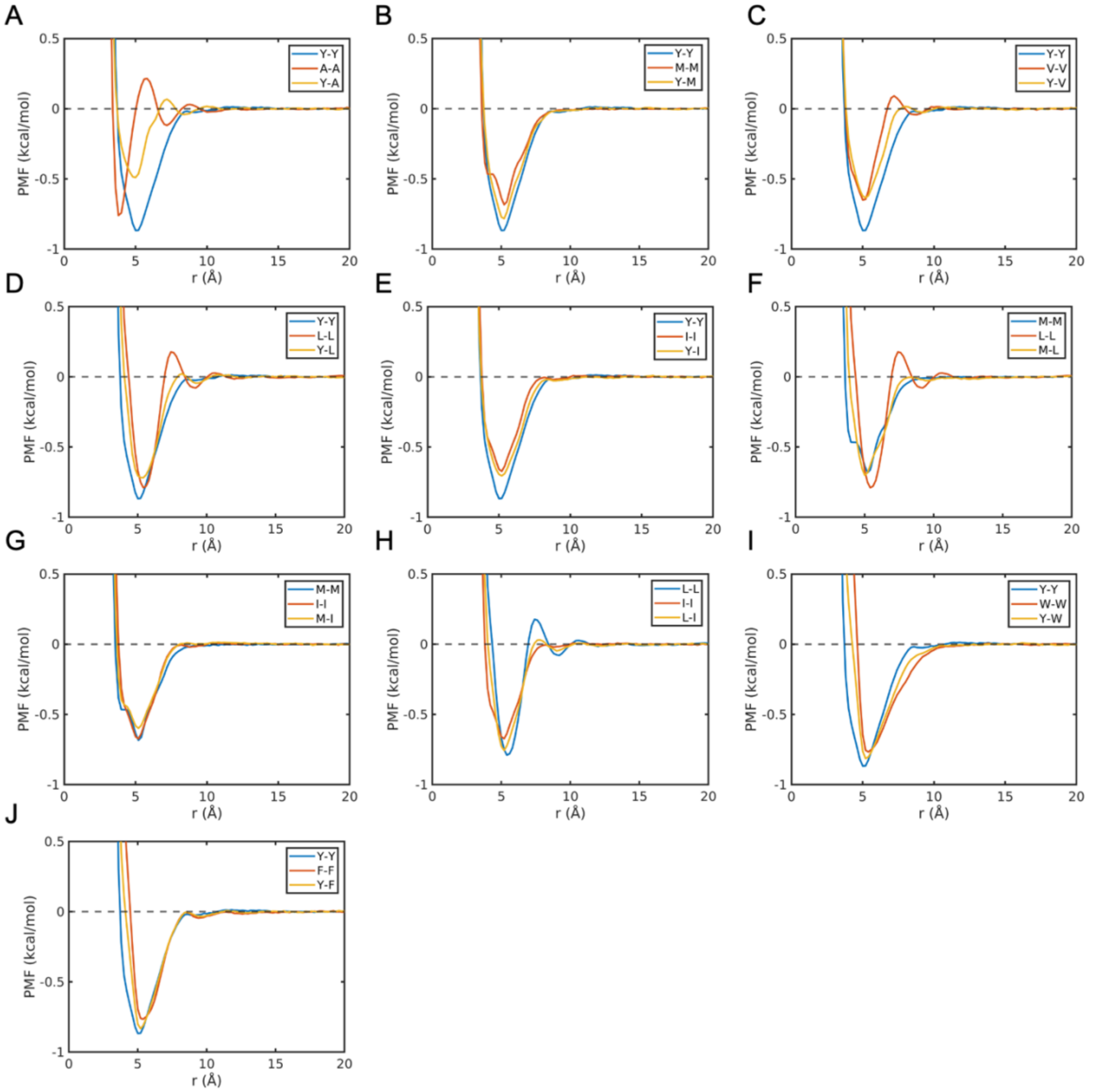


**Figure S6. PMFs for the ‘hydrophobic-hydrophobic’ heterotypic interactions at 300 K.** Heterotypic interactions are shown by yellow curves. For comparison, the corresponding homotypic interactions are shown in blue and red.

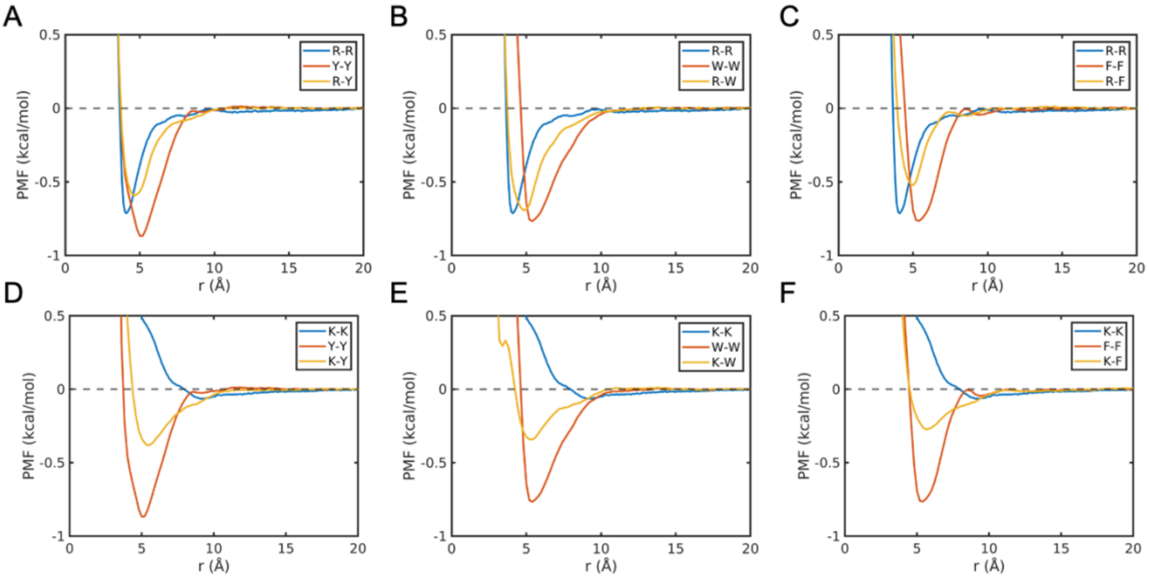


**Figure S7. PMFs for the 'cation-aromatic' heterotypic interactions at 300 K.** Heterotypic interactions are shown by yellow curves. For comparison, the corresponding homotypic interactions are shown in blue and red.

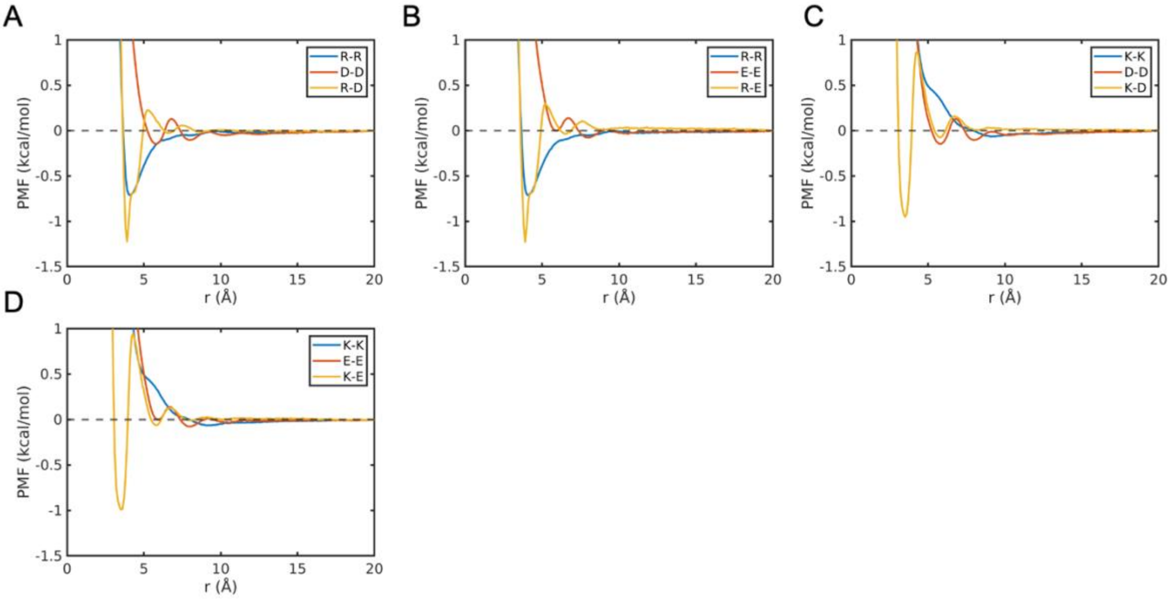


**Figure S8. PMFs for the 'charged-charged' heterotypic interactions at 300 K.** Heterotypic interactions are shown by yellow curves. For comparison, the corresponding homotypic interactions are shown in blue and red.

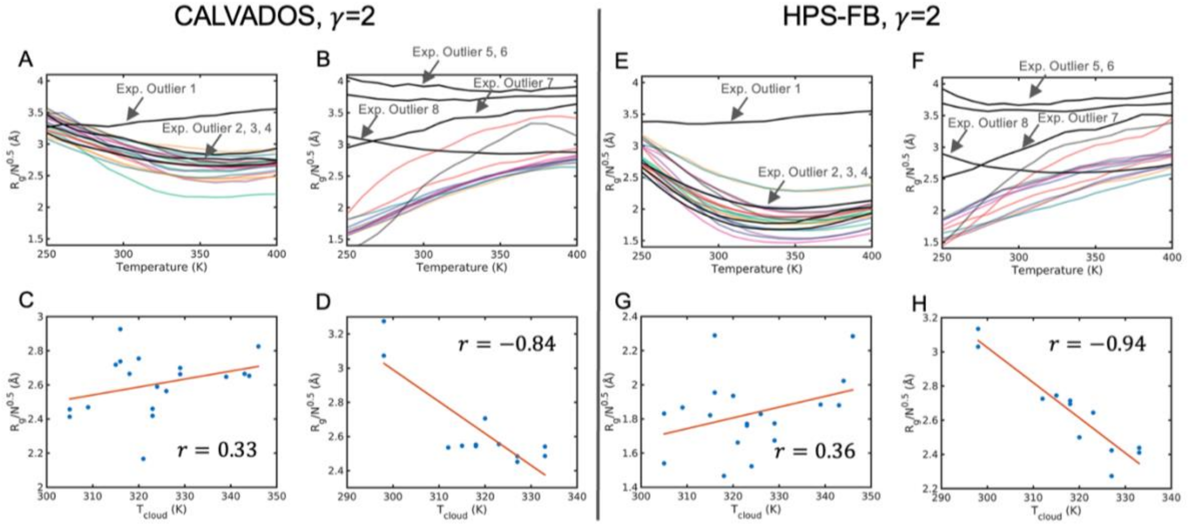


**Figure S9. Performance of TEA-augmented CALVADOS and HPS-FB models** ($\gamma = 2$). Comparison of simulated single-chain $R_g$ trends for ELP (LCST-type) and RLP (UCST-type) sequences. The models successfully distinguish between the two phase behaviors and identify experimental outliers (indicated by arrows).

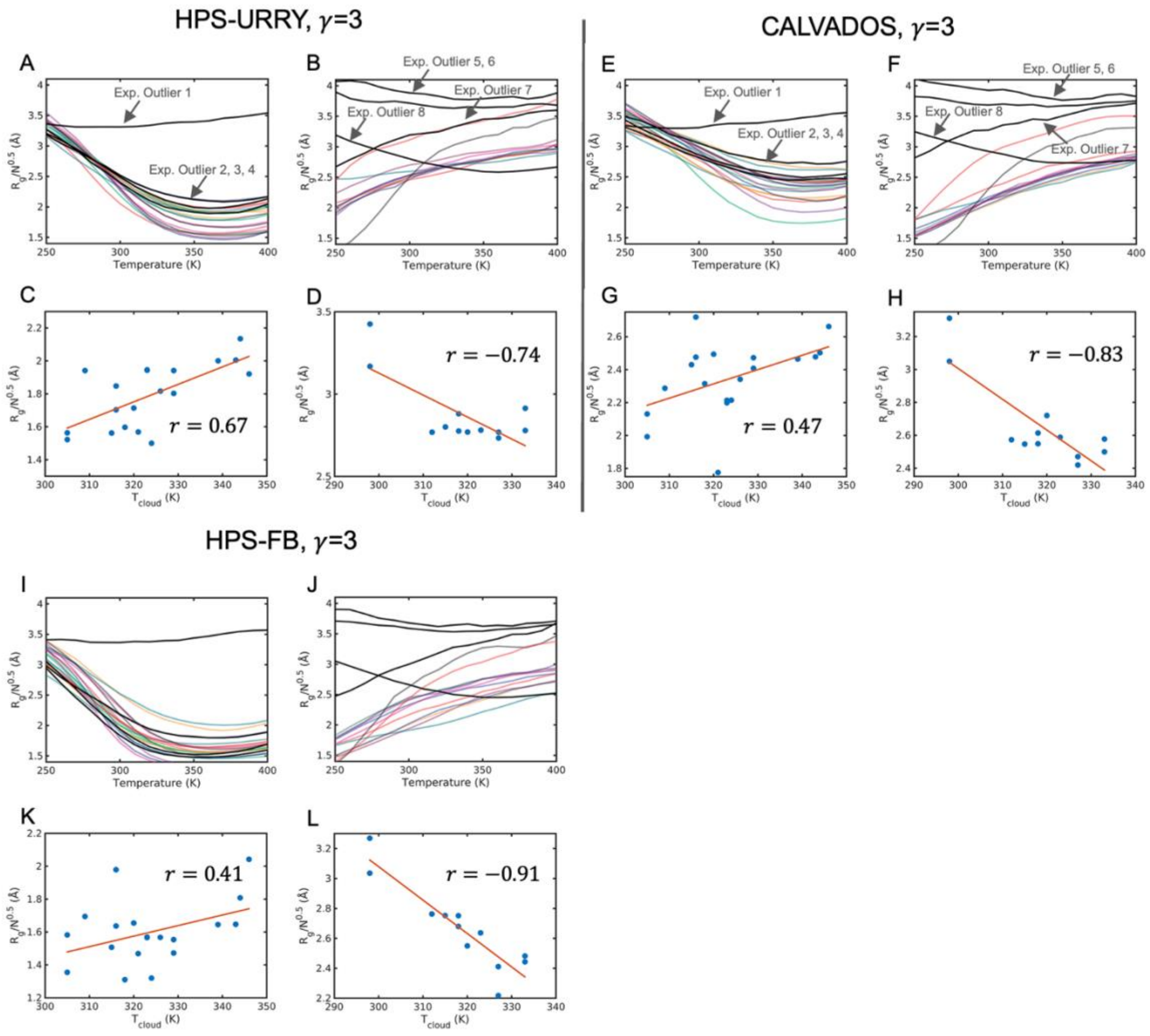


**Figure S10. Performance of TEA-augmented CALVADOS and HPS-FB models** ($\gamma = 3$). Comparison of simulated single-chain $R_g$ trends for ELP (LCST-type) and RLP (UCST-type) sequences. The models successfully distinguish between the two phase behaviors and identify experimental outliers (indicated by arrows).

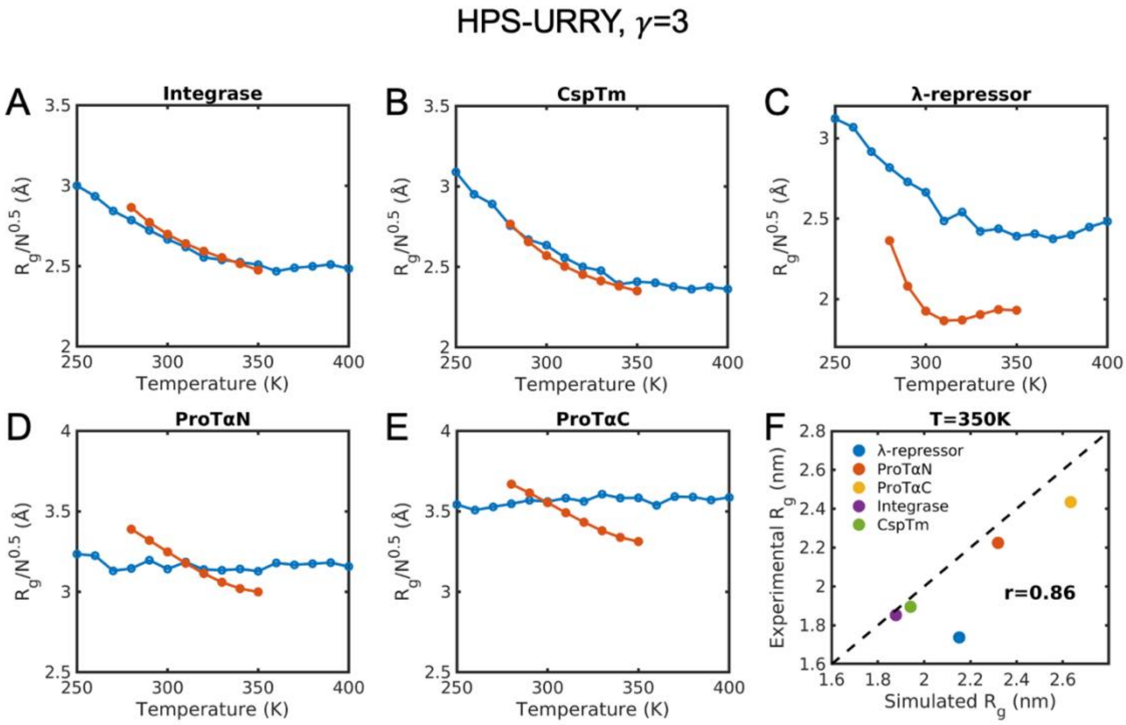


**Fig. S11. Comparison of simulated and experimental single-chain dimensions for five IDPs exihibiting LCST-type phase behavior.** Temperature-dpendent $R_g$ values from TEA-augmented HPS-Urry model with $\gamma = 3$ (blue curves) are compared with experimental measurements from Wuttke et al (red curves).

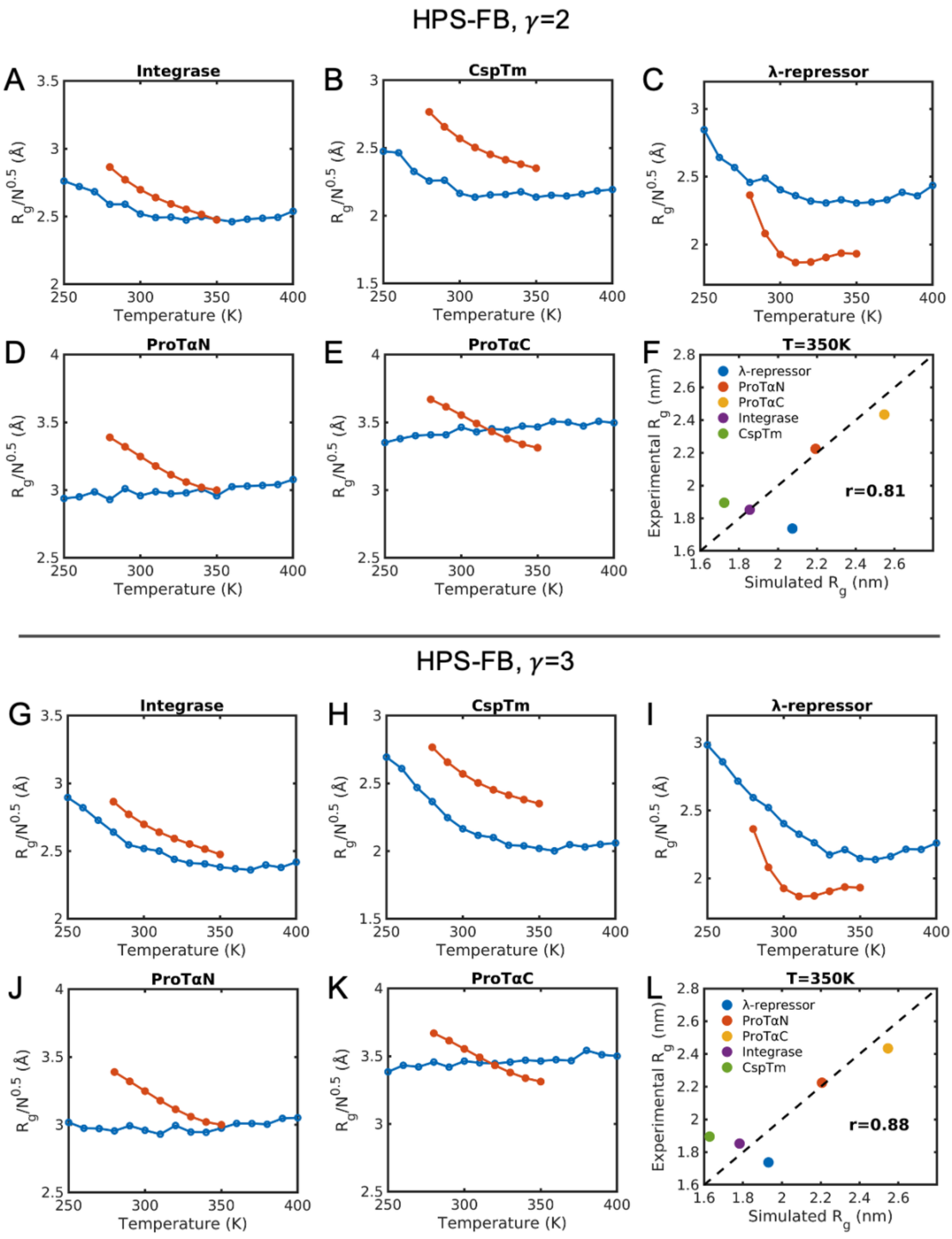


**Figure S12. Comparison of simulated and experimental single-chain dimensions for five IDPs exihibiting LCST-type phase behavior.** Temperature-dpendent $R_g$ values from TEA-augmented HPS-FB model (blue curves) are compared with experimental measurements from Wuttke et al (red curves).

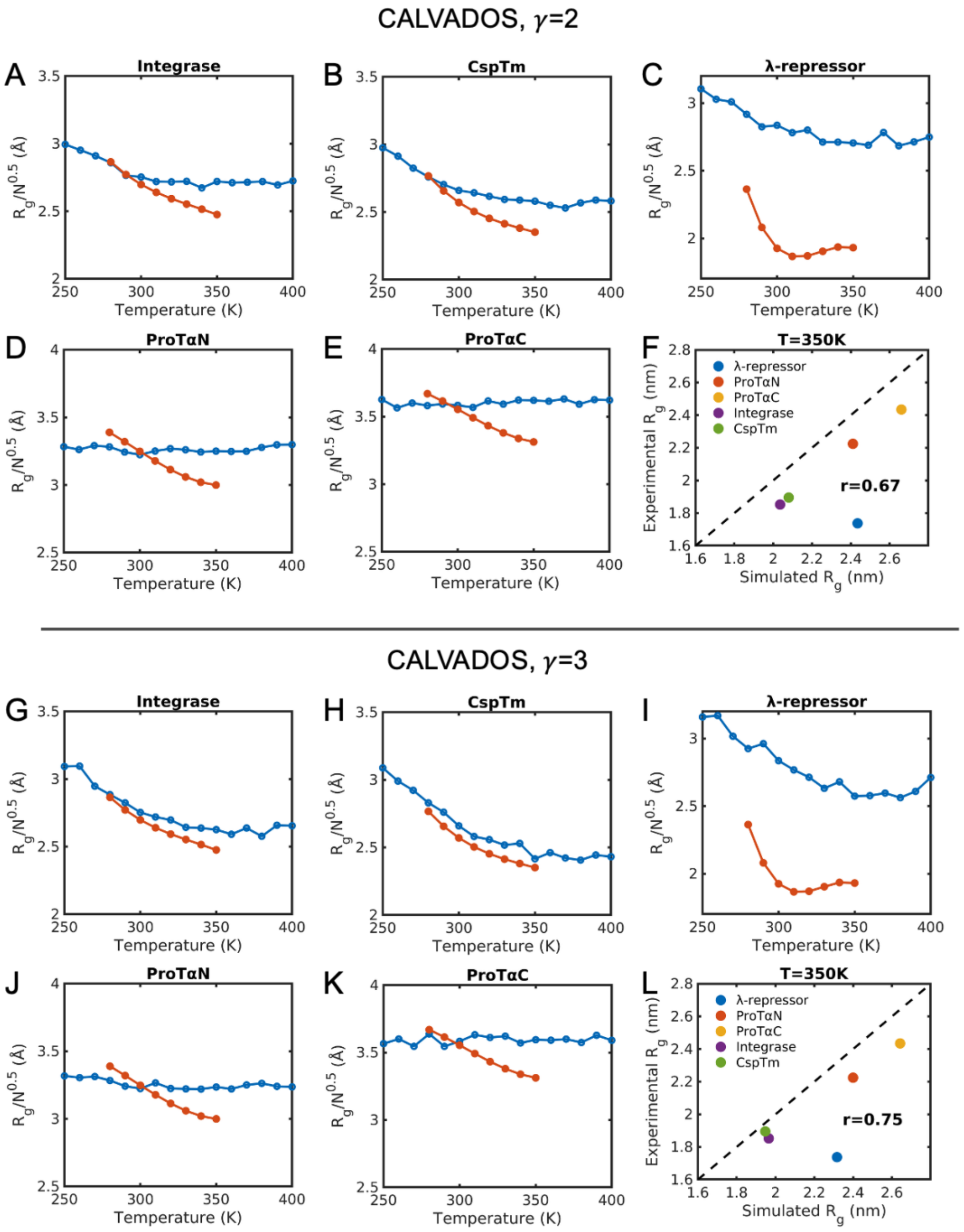


**Figure S13. Comparison of simulated and experimental single-chain dimensions for five IDPs exihibiting LCST-type phase behavior.** Temperature-dpendent $R_g$ values from TEA-augmented CALVADOS model (blue curves) are compared with experimental measurements from Wuttke et al (red curves)

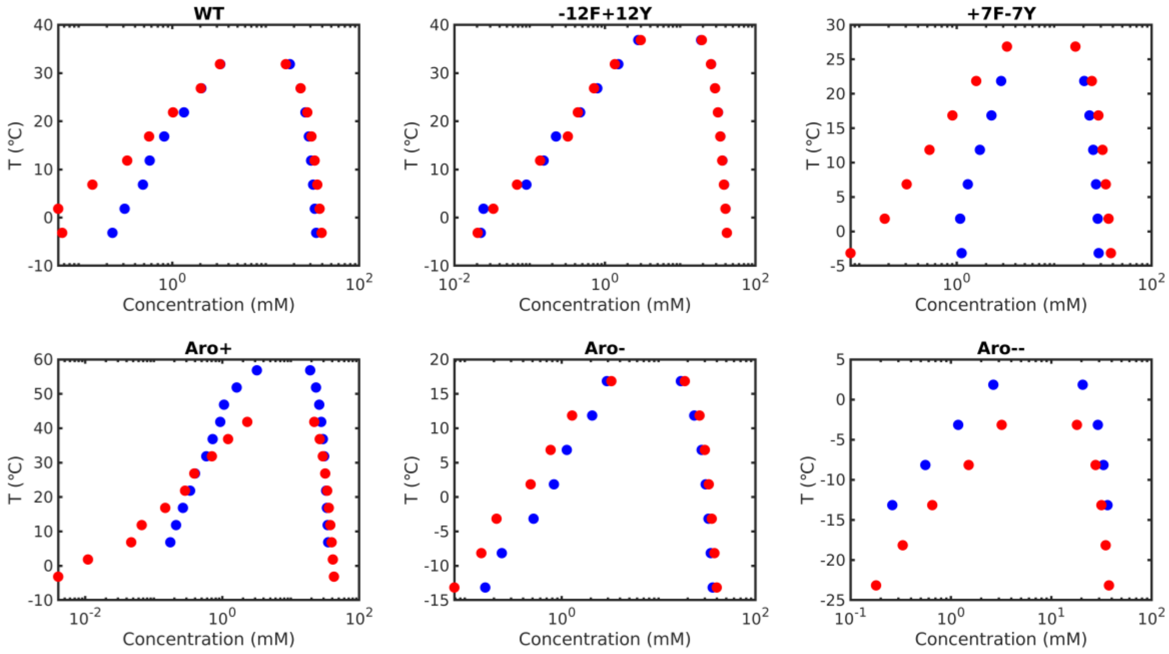


**Figure S14. Impact of temperature-dependent energetics on phase diagrams.** Comparison of phase diagrams from TEA-augmented HPS-Urry model (blue) versus the HPS-Urry model (red) for A1-LCD and its variants.

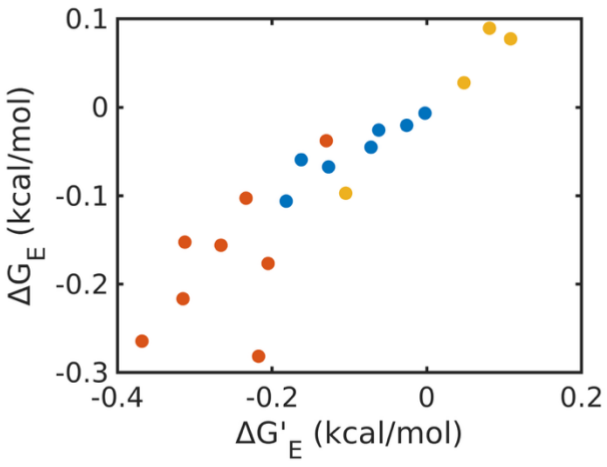


**Figure S15. Excess free energies calculated using the molecular-specific cutoff, $\Delta G'_E$, at 300 K correlates strongly with those derived using the 10 Å cutoff.**

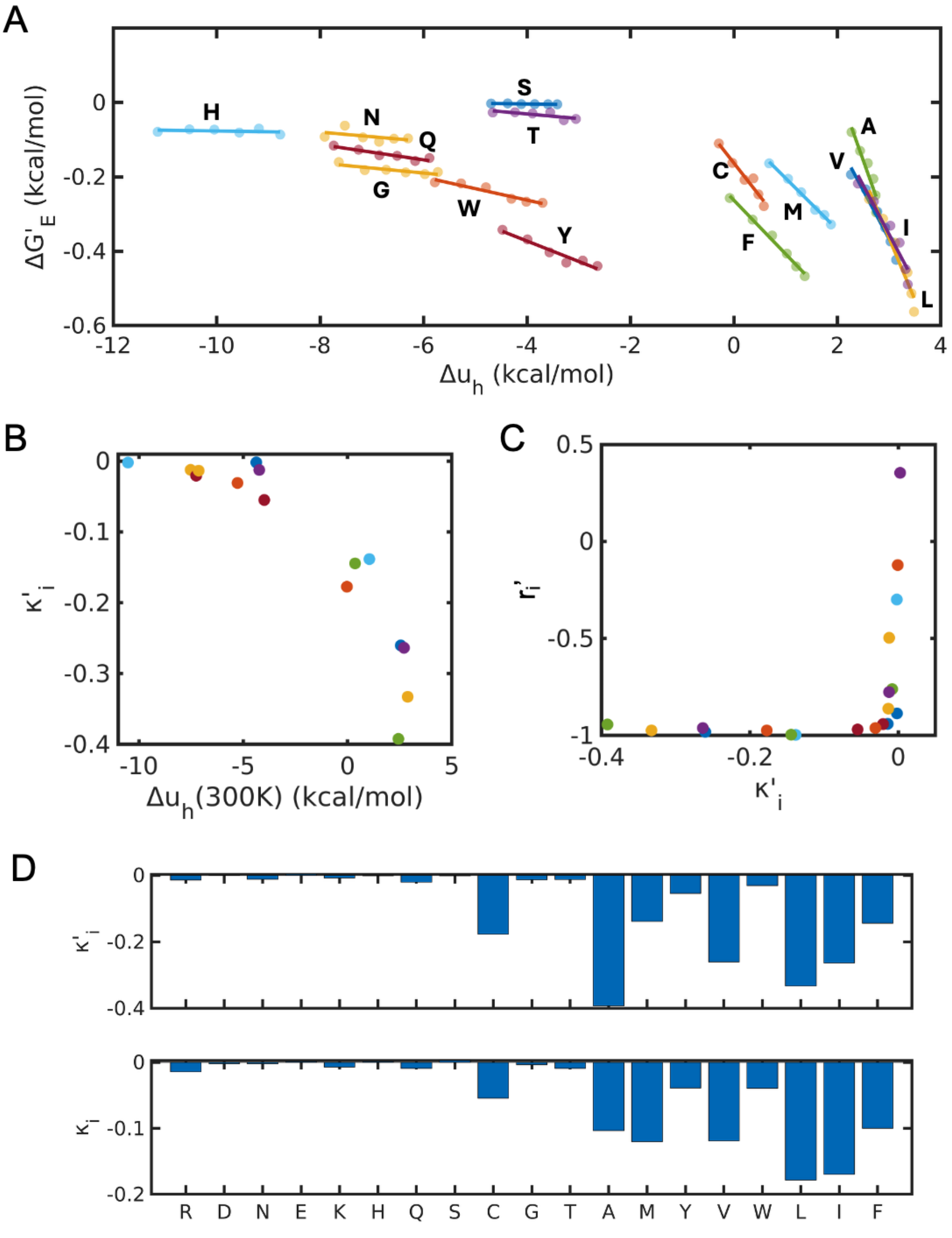


**Figure S16. Excess free energies $\Delta G'_E$ exhibit linear relationship with hydration free energies $\Delta u_h$.**(A) $\Delta G'_E$ versus $\Delta u_h$ plot for each analogue, (B) $\kappa'_i$ as a function of hydration free energies at 300 K $\Delta u_h(300K)$. (C) Dependence of the correlation coefficients $r'_i$ on the slope $\kappa_i$. (D) Comparison between $\kappa'_i$ and $\kappa_i$.

### Hydrophobic analogues

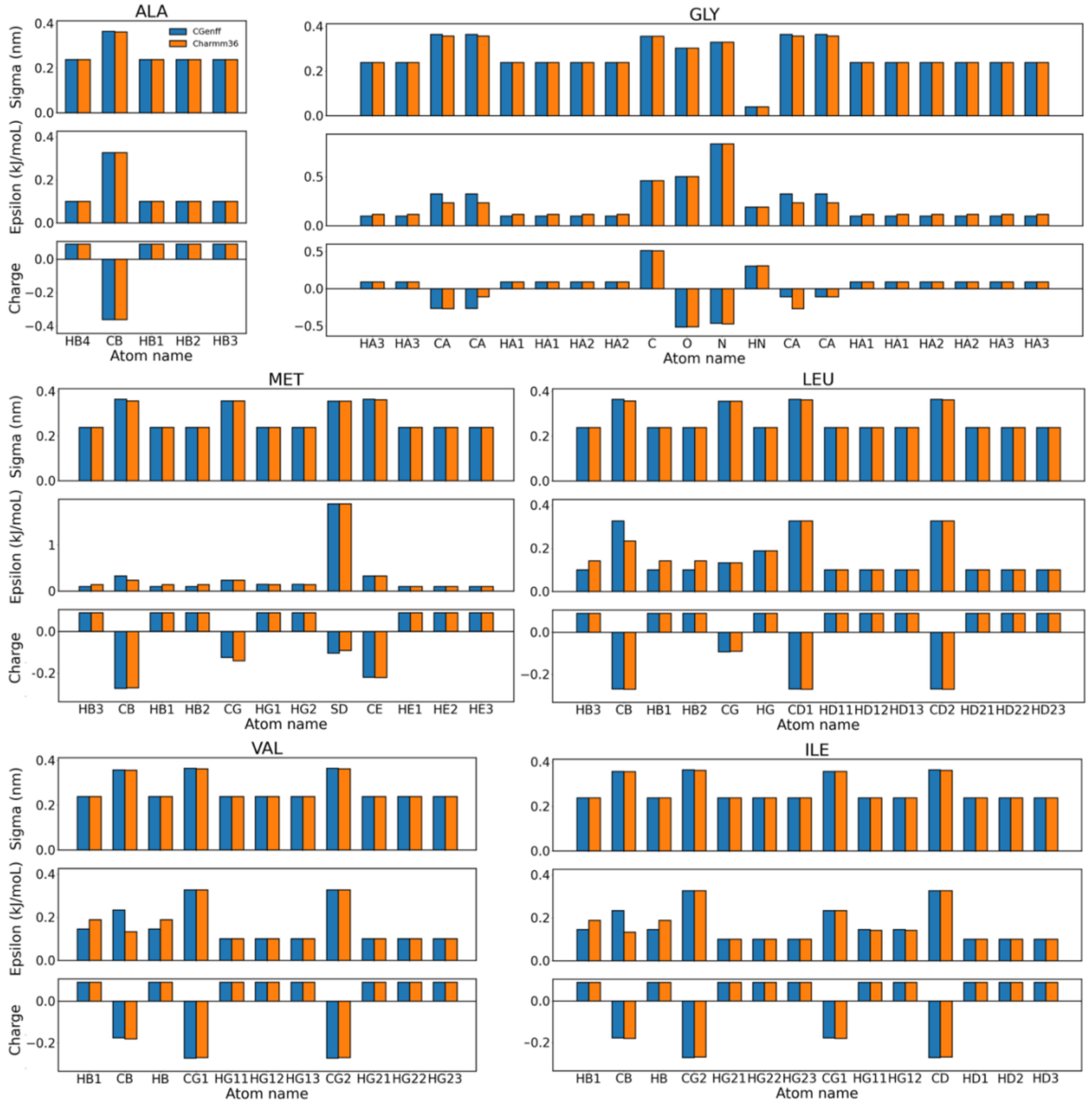


**Figure S17. Comparison of the sigma, epsilon, and partial charges in CGenFF (blue) and CHARMM36m protein force fields (red) for the hydrophobic analogues.**

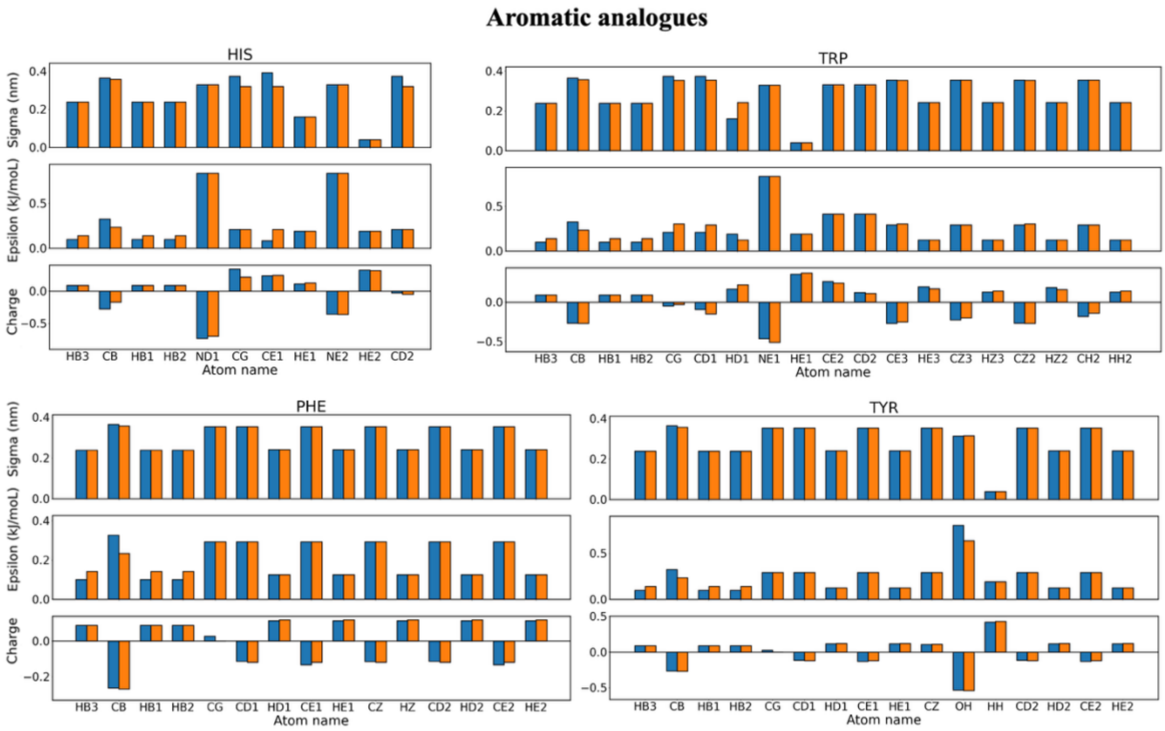


**Figure S18. Comparison of the sigma, epsilon, and partial charges in CGenFF (blue) and CHARMM36m protein force fields (red) for the aromatic analogues.**

#### Hydrophilic analogues

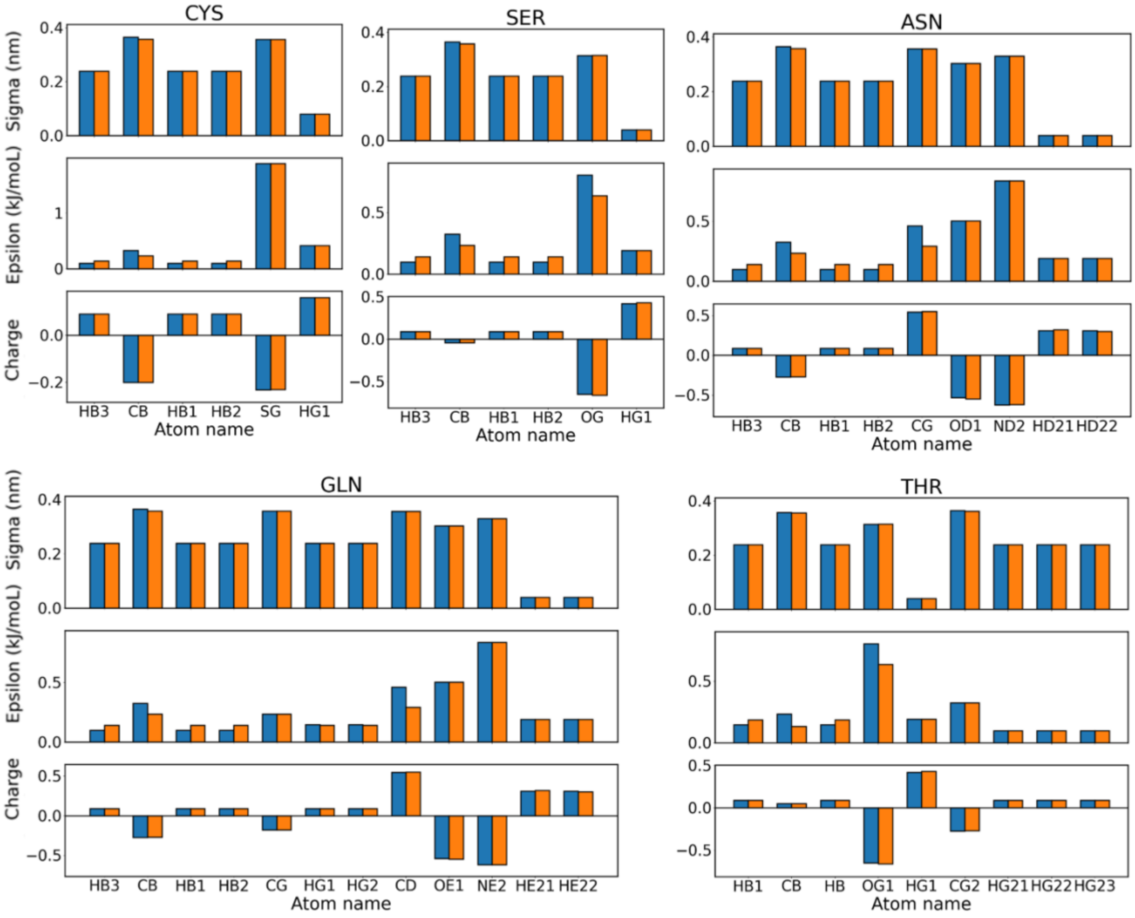


**Figure S19. Comparison of the sigma, epsilon, and partial charges in CGenFF (blue) and CHARMM36m protein force fields (red) for the hydrophilic analogues.**

#### Charged analogues

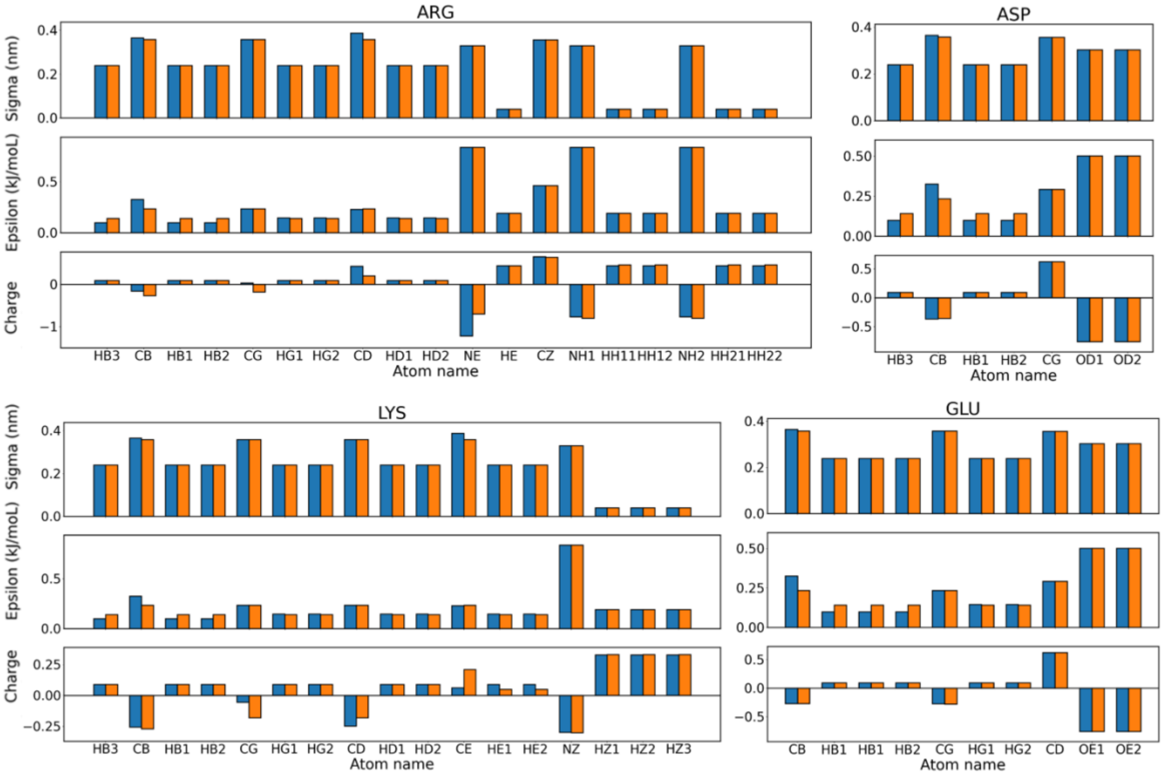


**Figure S20. Comparison of the sigma, epsilon, and partial charges in CGenFF (blue) and CHARMM36m protein force fields (red) for the charged analogues.**
